# Cell surface proteomics of patient-derived malignant rhabdoid tumor organoids identifies ROBO1 as potential CAR T cell target for pediatric solid tumors

**DOI:** 10.1101/2025.02.23.639405

**Authors:** Juliane L. Buhl, Jannet Koelewijn, Stijn De Munter, Victoria M. Cruz, Daryl Jun Kai Chin, Baojie Zhang, Marliek van Hoesel, Nienke van Herk, Irene Paassen, Katie Nachataya, Laura Hofman, Ella de Boed, Stefan Nierkens, Patrick Kemmeren, Annelisa M. Cornel, Wei Wu, Bart Vandekerckhove, Jarno Drost, Claudia Y. Janda

## Abstract

Malignant rhabdoid tumors are highly aggressive pediatric malignancies with limited treatment options and poor outcomes. To expand therapeutic options, we employed mass spectrometry-based cell surface proteomics on malignant rhabdoid tumor and patient-matched normal kidney organoids to identify potential CAR T cell targets. Integrating these findings with transcriptomics and protein expression data, we revealed ROBO1 as a promising target, showing strong and uniform expression across malignant rhabdoid and other pediatric tumors. ROBO1-targeted CAR T cells displayed potent anti-tumor activity *in vitro*, effectively eliminating tumor cells in co-cultures with organoids from malignant rhabdoid tumors, rhabdomyosarcoma, and neuroblastoma. *In vivo,* ROBO1 CAR T cells infiltrated tumors, induced potent tumor regression and significantly increased survival in malignant rhabdoid tumor-bearing mice. These findings establish ROBO1 as a compelling therapeutic target for CAR T cell therapy and offer a promising approach to address the critical need for effective treatments in high-risk pediatric solid tumors.

**Statement of significance:** This study identifies ROBO1 as a promising CAR T cell target for pediatric solid tumors. Using patient-derived tumor organoids and tissues, we demonstrate strong ROBO1 expression across several tumor entities and robust anti-tumor efficacy of ROBO1-targeted CAR T cells *in vitro* and *in vivo*, underscoring its potential for clinical translation.

## Introduction

Malignant rhabdoid tumors (MRTs) are rare and highly aggressive solid tumors that predominantly affect infants and young children. They arise in various anatomical sites, with extracranial malignant rhabdoid tumors (eMRTs) mostly occurring in the kidney and soft tissues, and atypical teratoid/rhabdoid tumors (ATRTs) developing in the central nervous system (CNS). MRTs typically develop during embryogenesis or early childhood, with most diagnoses occurring before the age of three (1–3). The treatment of MRTs is particularly challenging, as patients at this early age are highly susceptible to the toxic effects of chemotherapy and exhibit unique drug pharmacokinetics and pharmacodynamics due to immature organ systems. With current treatment – a combination of surgery, chemotherapy, and radiotherapy – the prognosis is dismal and the five-year overall survival rates are only around 20-30%, underscoring the urgent need for more effective therapeutic approaches (4).

MRTs are molecularly defined by the biallelic inactivation of *SMARCB1*, and less frequently *SMARCA4*. These genes encode core components of the SWI/SNF chromatin-remodeling complex, which plays a crucial role in regulating gene expression, and where absent, leads to profound epigenetic dysregulation (5–9). Beyond this hallmark alteration, MRTs exhibit a notably low tumor mutational burden and lack other oncogenic drivers (10,11). This complicates the development of effective conventional targeted therapies and neoantigen-dependent immunotherapies. However, advancements in molecular profiling and the development of preclinical models, such as patient-derived organoids that recapitulate the biology of patient tumors, are beginning to uncover MRT biology and open avenues for new therapeutic approaches (12–14).

Chimeric antigen receptor (CAR) T cell therapies have demonstrated remarkable success in treating hematologic malignancies, such as B cell leukemia and multiple myeloma, by targeting highly and uniformly expressed antigens like CD19 and BCMA (15). As the activity of CAR T cells does not depend on neoantigens, but rather on cell surface receptors, CAR T cells could offer promising therapeutic opportunities for MRTs. However, their efficacy against solid tumors remains limited due to challenges such as the scarcity of highly and uniformly expressed target antigens, an immunosuppressive tumor microenvironment, inefficient tumor infiltration, and limited CAR T cell persistence (16). Current basket trials enrolling MRT patients are investigating CAR T cell therapies targeting antigens such as the pan-pediatric cancer target B7-H3 (17,18), Glypican-3 (GPC3) (NCT06198296, NCT04377932, NCT04715191), and GD2 (NCT05298995, NCT03373097, NCT04099797). GPC3 and GD2 are predominantly expressed in a subset of pediatric solid tumors (*e.g*., liver tumors, neuroblastoma, bone sarcomas and brain tumors), but their variable expression in MRTs (19–21) limits trial accessibility to a subset of patients. While these trials show promise, most MRTs remain untreatable with CAR T cell therapies due to a lack of target expression.

Hence, expanding the repertoire of CAR T cell targets is critical to provide better therapeutic options for MRT patients. Claudin-6 (CLDN-6), a tight junction protein normally expressed during fetal development, has emerged as a promising CAR T cell target and is aberrantly upregulated in around 40% of MRTs (22). Preclinical studies and early clinical trials in adult patients have demonstrated encouraging results using CLDN-6-targeting CAR T cells in combination with mRNA vaccines designed to enhance CAR T cell persistence (23). Additionally, CAR T cells targeting CD70, a costimulatory receptor with limited expression in normal tissues (24), have shown efficacy in eliminating eMRT-derived organoids *in vitro* (25). However, CD70 expression has not been systematically evaluated in MRTs, and studies in a large patient cohort are needed to validate its expression and assess its therapeutic potential for MRTs.

Given the immense potential of CAR T cell therapy to improve outcomes for MRT patients, it is crucial to expand research efforts to identify uniformly expressed targets. Here, we used whole-organoid surface biotinylation, coupled to affinity purification and mass spectrometry to screen for CAR T cell targets in MRTs. By integrating proteomic and transcriptomic data from MRTs, other pediatric solid tumors, and normal tissues, we identified significant ROBO1 upregulation in MRTs and a wide range of other pediatric solid tumors, while exhibiting limited expression in postnatal normal tissues. ROBO1-targeted CAR T cells demonstrated robust activation, efficient tumor cell killing in co-culture with various pediatric tumor organoids *in vitro*, and complete tumor eradication in MRT patient-derived xenograft (PDX) models. These results highlight ROBO1 as a promising target for CAR T cell therapy, potentially addressing the scarcity of CAR T cell therapy options for pediatric MRTs and other solid tumors.

## Results

### Cell surface proteomics of patient-derived MRT organoids identifies potential targets for CAR T cell therapy

In this study, we leveraged patient-derived tumor and matched normal organoids previously developed in-house to identify and functionally evaluate novel CAR T cell targets for MRTs (12). Compared to conventional cell lines, tumor organoids offer a more accurate representation of the heterogeneity and maintain the genetic and transcriptional profiles of the original tumors (26,27). These features are particularly crucial for preclinical CAR T cell studies, as the therapeutic efficacy of CAR T cells heavily depends on the expression levels of cell surface antigens (28–30).

As a first step to identify potential targets for CAR T cell therapy, we analyzed the differential cell surface proteome of two patient-derived eMRT (of the kidney) organoid models (60T and 103T), and their patient-matched normal kidney organoids (60H and 103H) (12). Following cell surface biotinylation, cells were lysed, and biotinylated proteins were isolated, digested, and analyzed by mass spectrometry in triplicates (Fig. 1A-B). This analysis identified 1,971 biotinylated proteins, which were stringently filtered to yield 356 membrane-associated proteins (Suppl. Table 1). High correlation across the technical replicates demonstrated the experimental and technical robustness of this approach (Fig. S1A). Principal component analysis revealed that the normal organoid samples cluster together, underscoring their proteomic similarity, while the eMRT organoids from both patients showed distinct profiles of expressed membrane proteins, highlighting their unique cell surface proteomic signatures and dysregulated protein expression (Fig. S1B). Further analysis revealed that 48 and 73 proteins were significantly upregulated (fold change >2, p-value < 0.05) in 60T and 103T, respectively, with 118 and 100 proteins significantly downregulated by the same margin (Fig. 1C-D). Notably, 102 proteins were upregulated in either one or both eMRT models, of which 64 were categorized as receptors, 20 as kinases, 21 as transporters, and 12 as known cancer markers (Fig. S1C). These include CD70, an established MRT antigen (25), which showed significant overexpression in our analysis (Suppl. Table 1). However, CD276/B7-H3, another known MRT antigen (18), did not display notable differential expression between MRT and normal kidney organoids.

**Figure 1.**
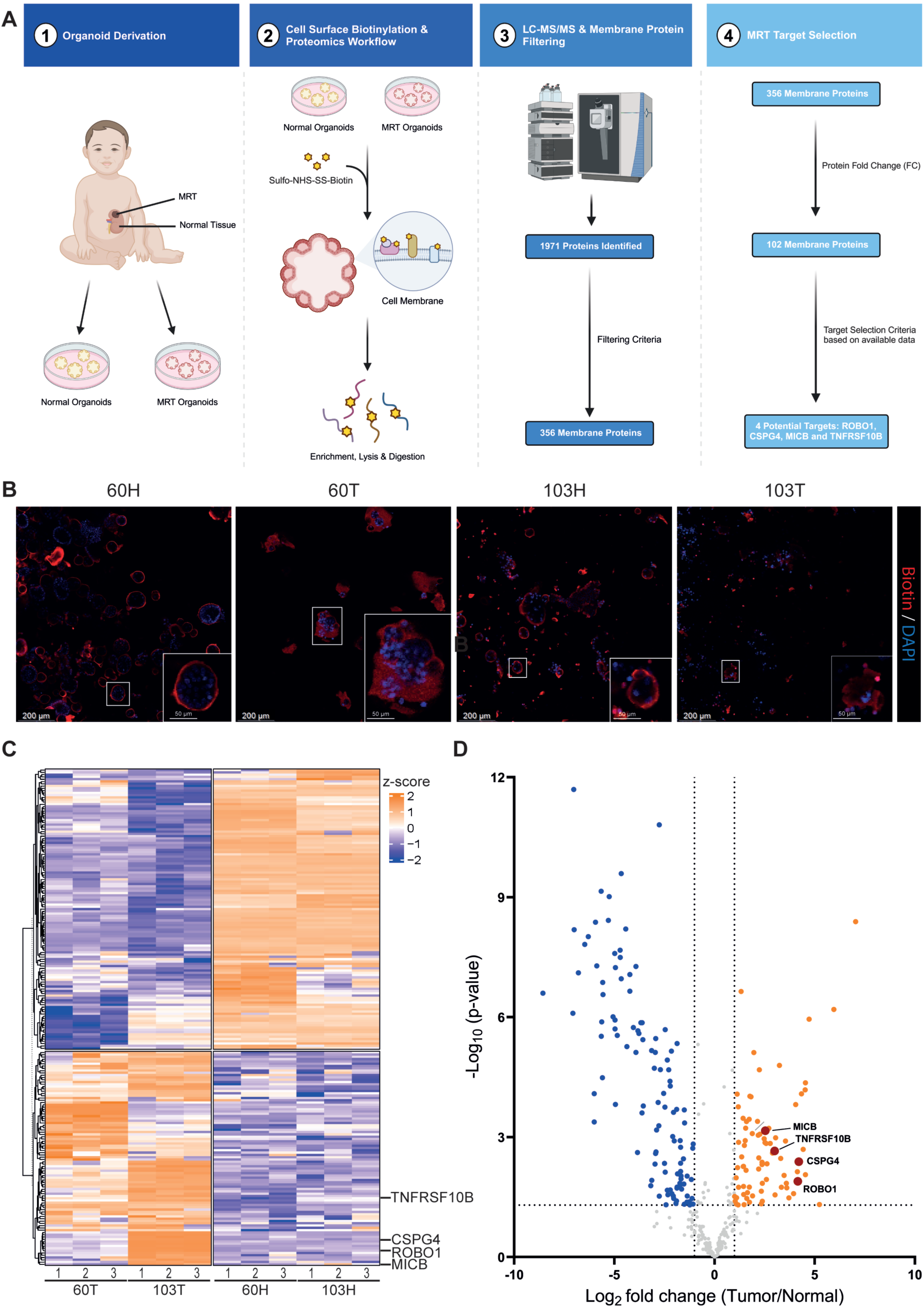
Cell surface proteomics of patient-derived MRT organoids identifies potential targets for CAR T cell therapy. (A) Schematics of the cell surface proteomics and data filtering workflow. (B) Immunofluorescence staining of biotinylated cell surface proteins (red) and cell nuclei (blue) in eMRT (T) organoids and patient-matched normal (H) kidney organoids. (C) Heatmap of significantly differentially expressed cell surface proteins in two paired eMRT and normal kidney organoid models. (D) Volcano plot illustrating differentially expressed cell surface proteins between eMRT and normal kidney organoids. Proteins with significant differential expression (fold change > 2, FDR-adjusted p-value < 0.05) are highlighted in blue and orange, while potential CAR T cell targets for eMRT are marked in red.

To refine potential CAR T cell targets in MRTs, we manually prioritized the membrane-associated proteins based on the following criteria (Fig. 1A): (1) high gene expression in MRTs (*i.e.*, both eMRTs and ATRTs) from public datasets (31,32), (2) relevance to cancer biology, (3) presence of both an extracellular and transmembrane domain, and (4) availability of antibodies tested in preclinical or clinical settings. This approach revealed four promising candidates (Fig. 1D): ROBO1 (Roundabout Guidance Receptor 1), which is crucial for neurodevelopment, particularly cell migration and axonal guidance, during embryonic development, and linked to tumor invasiveness when overexpressed (33,34); CSPG4 (Chondroitin Sulfate Proteoglycan 4), a cell surface proteoglycan associated with tumor progression and metastasis in various cancers (35,36); MICB (MHC class I polypeptide-related sequence B), a stress-induced receptor that activates immune responses and is frequently upregulated in tumors (37); and TNFRSF10B (also known as Death Receptor 5, DR5), a receptor for TRAIL-induced apoptosis that is commonly overexpressed in cancer cells, making it a target for inducing cell death in tumors (38,39).

### Gene and protein expression analysis validates ROBO1, CSPG4, MICB and TNFRSF10B upregulation in MRT organoids

To validate our mass spectrometry findings and support the selection of ROBO1, CSPG4, MICB, and TNFRSF10B as potential CAR T cell targets in MRTs, we next evaluated their RNA and protein expression in the 60T and 103T eMRT organoid models that were used for the cell surface proteomics experiments, as well as additional eMRT organoid models. Using available bulk RNA sequencing (RNA-seq) data (12), flow cytometry, and Western blotting, we compared their expression levels to those in normal kidney organoids (Fig. 2A-B, S2A).

**Figure 2.**
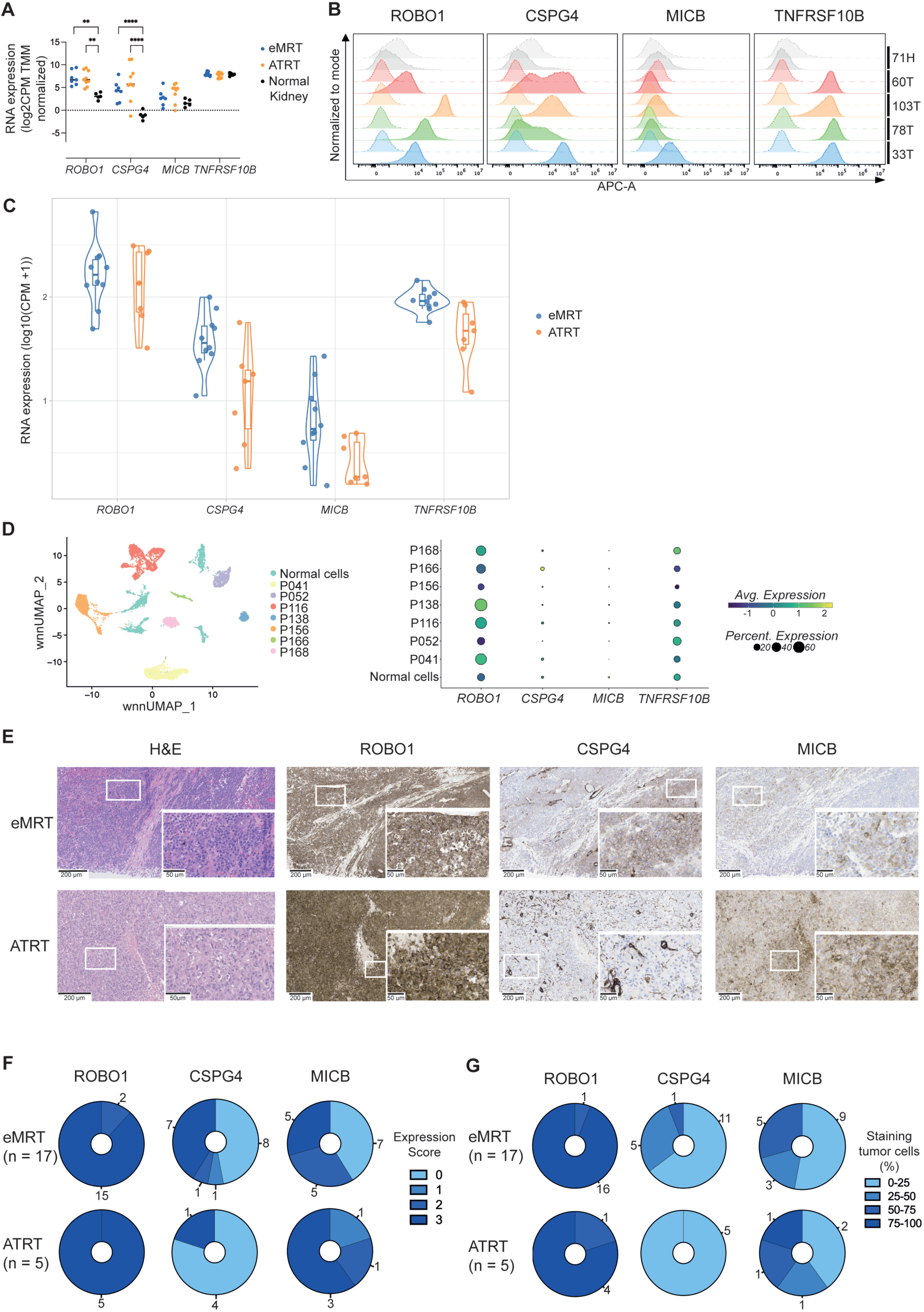
Gene and protein expression analysis validates ROBO1, CSPG4, MICB and TNFRSF10B upregulation in MRT organoids and tissues. (A) RNA expression levels of *ROBO1*, *CSPG4*, *MICB,* and *TNFRSF10B* in normal kidney (n=5), eMRT (n=7), and ATRT (n=9) organoids. Data are represented as individual values with the median indicated by a line. Statistical significance is represented as follows: *p < 0.05, **p < 0.01, ***p < 0.001, ****p < 0.0001. (B) Flow cytometry histogram plots showing surface expression of ROBO1, CSPG4, MICB, and TNFRSF10B expression in one normal kidney organoid (H) model and four eMRT (T) organoid models. Dashed lines indicate isotype controls. Representative data from at least two biological replicates. (C) Violin plots depicting *ROBO1, CSPG4, MICB, and TNFRSF10B* RNA expression levels in eMRT (n=10) and ATRT (n=7) tissues. (D) UMAP visualization of normal cells clusters as well as tumor cell clusters that segregate by patient (P) in eMRT tissues, with a dot plot showing *ROBO1*, *CSPG4*, *MICB,* and *TNFRSF10B* expression across samples. (E) Representative IHC images of ROBO1, CSPG4, and MICB expression in eMRT (n=17) and ATRT (n=5) tissues, with quantification of expression scores (F) and fraction of positive cells (G).

Our proteomics data indicated high ROBO1 overexpression in 103T and moderate overexpression in 60T (Fig. S1D). This aligned with bulk RNA-seq results, which showed elevated *ROBO1* mRNA expression in eMRT organoids compared to normal kidney organoids (Fig. 2A). Flow cytometry confirmed robust ROBO1 cell surface protein expression across eMRT organoid models compared to normal kidney organoids (Fig. 2B), while Western blotting, which is less sensitive than flow cytometry and detects total protein expression, detected ROBO1 in the highest expressing lines 103T and 33T (Fig. S2A). CSPG4 showed a similar expression pattern to ROBO1, with strong expression in 103T and moderate expression in 60T by proteomics. CSPG4 RNA and protein levels were elevated across most eMRT organoid models compared to normal kidney organoids, with 78T exhibiting somewhat lower expression (Fig. 2A-B, S2A). Yet, CSPG4 protein expression varied both between and within organoid models.

In contrast to ROBO1 and CSPG4, MICB was marginally overexpressed in 103T and 60T based on proteomics data, which aligns with results from bulk RNA-seq, flow cytometry and Western blotting (Fig. 2A-B, S2A). In addition, TNFRSF10B was moderately overexpressed in 103T and 60T according to proteomics. Although RNA expression was not elevated in eMRT organoid models relative to normal kidney organoids, protein levels were significantly higher as confirmed by both flow cytometry and Western blot (Fig. 2A-B, S2A). This discrepancy suggests post-translational regulation of TNFRSF10B expression in non-malignant cells and shows the added value of target discovery using proteomics approaches (40). Furthermore, analysis of existing bulk RNA-seq data from ATRT and MRT brain metastasis (BM)-derived organoids (14), revealed that RNA expression followed similar trends to those observed in eMRTs (Fig. 2A). Collectively, these results confirm the overexpression of ROBO1, CSPG4, and TNFRSF10B, and comparable expression of MICB in MRT organoids relative to normal kidney organoids.

### ROBO1 is strongly and uniformly expressed in patient-derived eMRT and ATRT tissues

High and uniform expression of CAR T cell targets within and across tumors is critical for maximizing therapeutic efficacy and broadening their clinical application. However, most CAR T cell targets in solid tumors exhibit heterogeneous expression, posing major challenges. Intratumoral heterogeneity promotes tumor progression by enabling the survival and proliferation of antigen-negative or antigen-low clones, ultimately leading to resistance against CAR T cell therapy (28–30,41). To address this, we evaluated both intra- and intertumoral heterogeneity of the four shortlisted markers in eMRT and ATRT tissues.

We first analyzed bulk RNA-seq data from eMRT (n=10) and ATRT (n=7) tumors treated at our center (42). Consistent with RNA-seq data from the organoids, *ROBO1* was strongly expressed in both eMRTs and ATRTs, while *CSPG4* and *TNFRSF10B* showed intermediate expression levels, and *MICB* showed only weak expression (Fig. 2C). To assess intratumoral expression heterogeneity, we evaluated RNA expression of the four targets using single-nuclei RNA sequencing data from seven eMRT tissue samples generated previously in-house (Fig. 2D, S2B) (9). Notably, *ROBO1* RNA was expressed in the majority of tumor cells and samples (5 out of 7). In contrast, *TNFRSF10B* was expressed in a smaller subset, while *CSPG4* and *MICB* were expressed at a low level in only a fraction of tumor cells.

To validate these findings at the protein level, we performed immunohistochemistry (IHC) on paraffin-embedded tumor sections from eMRT (n=17) and ATRT (n=5) samples (Fig. 2E-G, Suppl. Table 2), using marker-specific antibodies. Of note, validation for TNFRSF10B could not be performed due to the unavailability of a suitable antibody. Consistent with our RNA expression analyses, ROBO1 was strongly and uniformly expressed across all eMRTs and ATRT samples, with 20 of 22 samples scoring the highest expression level (score 3) and 75-100% of tumor cells expressing ROBO1. In contrast, MICB and CSPG4 exhibited limited and heterogenous expression, with non-uniform staining patterns within individual samples. In summary, while ROBO1 is strongly and consistently expressed in MRTs, CSPG4, TNFRSF10B and MICB exhibit variable expression, making these antigens less favorable targets for CAR T cell therapy.

Low expression in normal tissues is crucial to minimize on-target, off-tumor toxicity (43). Therefore, we next evaluated ROBO1 expression in normal tissues at both the RNA and protein levels. Given that MRTs typically develop during embryogenesis and are generally diagnosed before the age of three, we reverted to a publicly available transcriptomic dataset that includes seven organs across development stages, from early organogenesis to adulthood (44). *ROBO1* RNA expression was high in the brain/cerebellum, heart, kidney and liver during embryonal development but declined markedly during the fetal stage and was largely downregulated after birth (Fig. S2C), consistent with its well-established role in neurodevelopment (33,34). We further examined ROBO1 protein expression in normal tissues commonly affected by MRTs, including the brain, kidney and liver, by IHC (Fig. S2D). ROBO1 was only weakly expressed in critical brain regions, such as the cerebellum, frontal lobe, and pons, and normal liver tissue. Similarly, kidney tissues, including the cortex and medulla, showed little to no ROBO1 expression, with only some expression detected in the glomeruli. Taken together, ROBO1 emerges as the most promising CAR T cell target due to its strong and uniform expression across MRTs and its minimal expression in essential normal tissues, supporting its further preclinical evaluation.

### ROBO1-targeting CAR T cells elicit strong and specific anti-tumor activity in eMRT organoids

To functionally evaluate ROBO1 as a target for CAR T cell therapy in MRTs, we performed *in vitro* co-culture experiments with eMRT organoids, assessing CAR T cell activation, cytokine release, and anti-tumor efficacy. ROBO1-targeting CAR T cells were generated using a previously described ROBO1-binding single-chain variable fragment (scFv) (Fig. S3A-B) (45), incorporated into a second-generation CAR lentiviral vector, encoding the CD8a hinge and transmembrane region, and 4-1BB/CD3ζ co-stimulatory domains (Fig. 3A). A CD19-targeting CAR served as a control, as CD19 is absent in MRTs. CAR T cells with either specificities were generated from healthy donor-derived T cells, followed by sorting the CAR-positive T cell population, and expansion (Fig. S3C).

**Figure 3.**
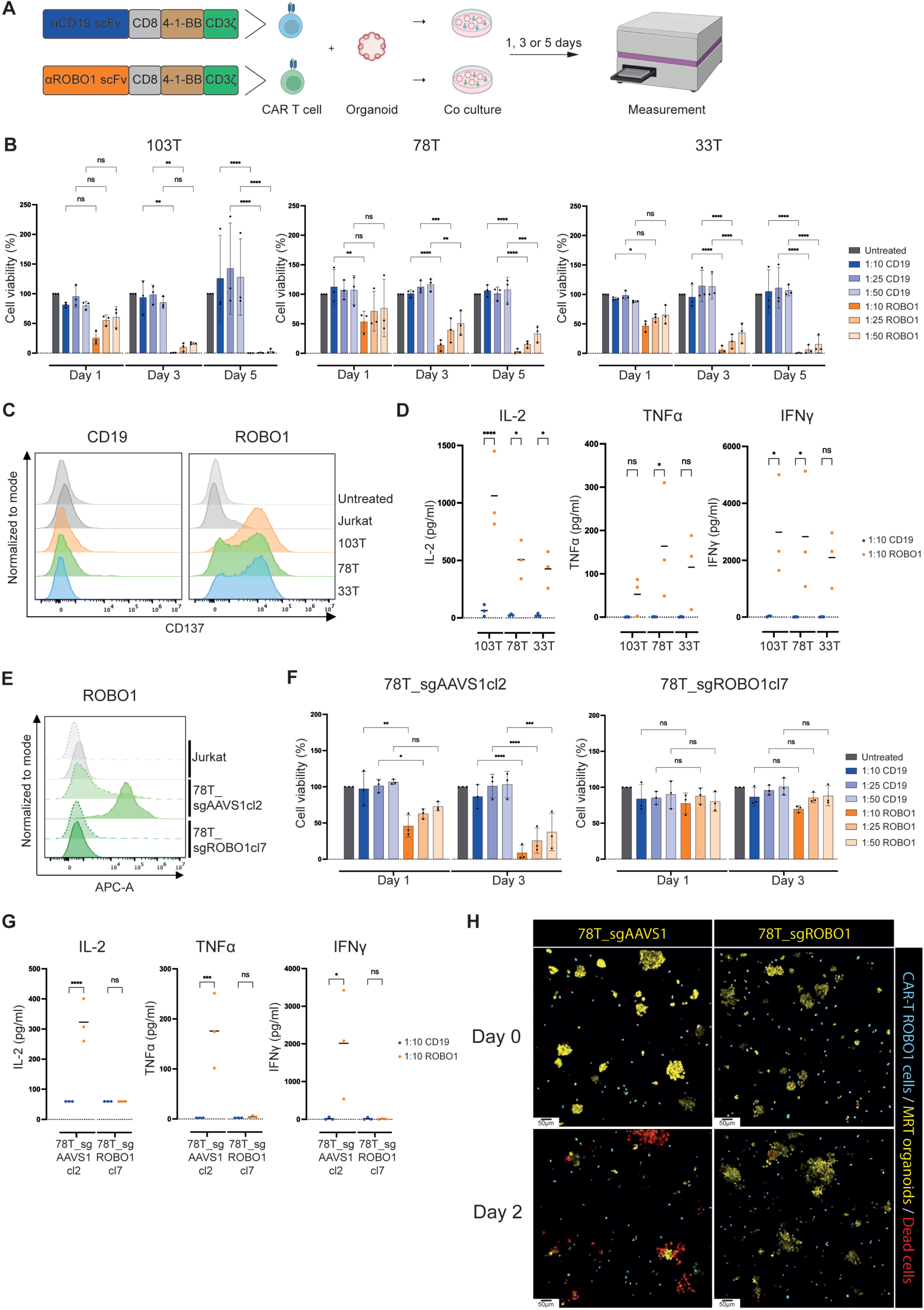
ROBO1-targeting CAR T cells elicit strong and specific anti-tumor activity in eMRT organoids. (A) Schematic of ROBO1 and CD19 CAR constructs and cell viability assay workflow. (B) Cell viability of eMRT organoids acer co-culture with ROBO1/CD19 CAR T cells at different E:T ratios. Results were normalized to untreated organoids. (C) Flow cytometry histogram plots of CD137 expression on ROBO1/CD19 CAR T cells acer 20-hour co-cultures with eMRT organoids, ROBO1-negative Jurkat cells or untreated at an E:T of 1:10. (D) Cytokine production acer 1-day co-culture of CAR T cells with eMRT organoids at an E:T of 1:10. (E) Flow cytometry histogram plots of ROBO1 expression on Jurkat cells, control (78T_sgAAVS1cl2) or ROBO1 (78T_sgROBO1cl7) knock-out eMRT organoids. Dashed lines indicate staining with isotype control antibody. (F) Cell viability of control or ROBO1 knock-out eMRT organoids acer co-culture with CAR T cells. Results were normalized to untreated organoids. (G) Cytokine production acer 1-day co-culture of CAR T cells with control or ROBO1 knock-out eMRT organoids. (H) 3D immunofluorescence imaging of ROBO1 CAR T cells co-cultured with control or ROBO1 knock-out eMRT organoids on day 0 and 2 at an E:T ratio of 1:10. Maximum intensity projections shown. Scale bars = 50 µm. Data in (B) and (F) represent the mean ± SD of three biological replicates, each containing the mean of three technical replicates, and data in (D) and (G) represent the mean of three biological replicates, each containing the mean of two technical replicates. (C) and (E) show representative data from three biological replicates. Statistical significance as in Fig. 2A.

Co-cultures were set up with ROBO1 and CD19 CAR T cells and firefly luciferase-expressing eMRT organoids (103T, 78T and 33T), selected based on their moderate to high ROBO1 cell surface expression (Fig. 2B). CAR T cells were added at effector-to-target (E:T) cell ratios of 1:50, 1:25 and 1:10, and tumor organoid viability was assessed on day 1, 3 and 5 using luminescence as a readout. ROBO1 CAR T cells exhibited dose-dependent cytotoxicity across all eMRT models compared to the CD19 CAR T cells (Fig. 3B). Notably, the highly ROBO1-expressing 103T organoids were nearly eradicated by day 5 at the lowest E:T ratio (1:50), while the moderately expressing 78T and 33T lines required higher ratios (1:10) for comparable cytotoxicity, consistent with an antigen density-depended response (28–30). Tumor cell killing by ROBO1 CAR T cells was associated with upregulation of CD137, a marker of T cell activation, as assessed by flow cytometry (Fig. 3C), and the secretion of pro-inflammatory cytokines (IL-2, TNF-α, and IFN-γ) (Fig. 3D, S3D). In contrast, CD19 CAR T cells showed minimal cytotoxicity, T cell activation and cytokine release (Fig. 3B-D, S3D). To further support ROBO1-dependency of the observed CAR T cell cytotoxicity, we generated clonal ROBO1 (sgROBO1) and control (sgAAVS1) knock-out tumor organoids from the 78T line using CRISPR-Cas9 (Fig. 3E, S3E-F). ROBO1 CAR T cells failed to kill sgROBO1 78T organoids and showed baseline cytokine release (Fig. 3F-G, S3G-H), while retaining robust cytotoxicity against sgAAVS1 (ROBO1 wild-type) 78T organoids. Furthermore, live-cell imaging confirmed selective cytotoxicity toward sgAAVS1 organoids, confirming the requirement of ROBO1 expression for the observed cytotoxicity of ROBO1-targeting CAR T cells (Fig. 3H).

### ROBO1 is a promising CAR T cell target for other pediatric solid tumors

To evaluate the broader applicability of ROBO1 as CAR T cell target in pediatric oncology, we first analyzed *ROBO1* RNA expression across a range of pediatric solid and brain tumors (Fig. 4A-B) (42). Bulk RNA sequencing data from 372 pediatric tumor samples, spanning renal, hepatic, bone, soft tissue, and CNS tumors revealed high *ROBO1* expression in several tumor types, particularly in rhabdomyosarcoma (RMS), neuroblastoma (NB), Wilms tumor (WT), osteosarcoma (OS), glioblastoma (GBM), and hepatoblastoma (HB), in addition to eMRTs and ATRTs. Notably, *ROBO1* expression remained robust across different patient age groups (Fig. S4A). Furthermore, although already deprioritized, *CSPG4*, *MICB*, and *TNFRSR10B* show a more tumor subtype-specific expression profile compared to *ROBO1* (Fig. S4A). A recent study reported *ROBO1* overexpression in a patient-derived tumor line from pediatric medulloblastoma (classified as subgroup 3, one of the four consensus molecular subgroups) (46). In our transcriptomic analysis, *ROBO1* expression in medulloblastoma was high, though patient-to-patient variability was observed (Fig. 4B). This variability appeared to correlate with the different medulloblastoma subgroups, showing higher expression in the WNT and SHH subgroups, and heterogenous expression in group 3 and 4, consistent with previous findings (46) (Fig. S4B). Furthermore, IHC on pediatric tumor tissues confirmed strong, uniform ROBO1 expression across most tested tumors (*e.g.,* NB, medulloblastoma (MB), WT, and renal cell carcinoma (RCC)) and variable expression in RMS and Ewing sarcoma (ES) (Fig. 4C-D, S4C-D).

**Figure 4.**
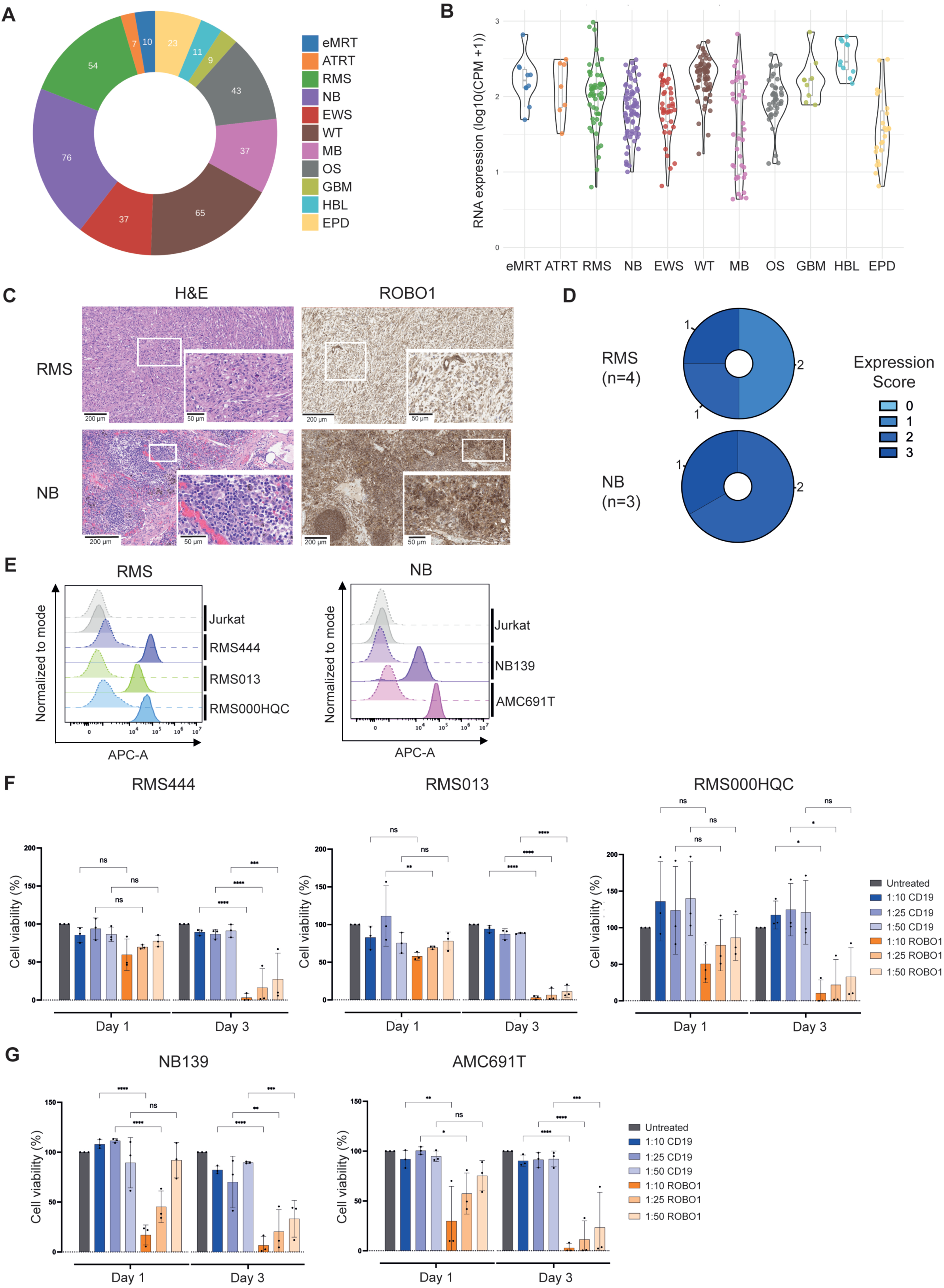
ROBO1 is a promising CAR T cell target for other pediatric solid tumors. (A) Pie chart showing the composition of the patient cohort used for RNA-seq analysis in (B). (B) Violin plot of *ROBO1* RNA expression across pediatric solid and brain tumors. (C) Representative IHC images of ROBO1 expression in RMS (n=4) and NB (n=3) tumor tissues, with quantification of expression scores and fraction of positive cells in (D). (E) Flow cytometry histogram plots of ROBO1 expression on Jurkat cells, three RMS and two NB organoid models. Dashed lines indicate isotype controls. Representative data from three biological replicates. (F) Cell viability of three RMS and two NB organoid models (G) acer co-culture with ROBO1/CD19 CAR T cells at different E:T ratios. Data represent mean ± SD of three biological replicates, each containing the mean of at least three technical replicates. Statistical significance as in Fig. 2A.

Next, we evaluated the anti-tumor activity of ROBO1-targeting CAR T cells in NB and RMS, given the availability of patient-derived tumor organoids. Co-culture experiments were performed with three independent patient-derived RMS organoid models (RMS444, RMS013, RMS000HQC) (47) and two NB organoid models (NB139, AMC691T) (48), representing diverse molecular subtypes, including fusion and MYCN amplification status, respectively. Expression analysis confirmed high ROBO1 cell surface expression in RMS as well as in NB organoids (Fig. 4E). ROBO1-targeting CAR T cells demonstrated potent cytotoxicity across all tested tumor organoid models compared to CD19-targeting CAR T cells. At an E:T ratio of 1:10, nearly complete tumor cell eradication was achieved by day 3, while even at the lowest E:T ratio of 1:50, over 50% tumor cell killing was observed (Fig. 4F-G).

### ROBO1-targeting CAR T cells induce potent and sustained cytotoxicity, eradicating eMRTs in mice

To preclinically evaluate the potential of ROBO1 as a CAR T cell target *in vivo*, we used MRT PDX models. We first conducted a dose-finding study followed by efficacy experiments. Immunodeficient mice were subcutaneously injected with luciferase-expressing 33T and 103T MRT organoids, and tumors were allowed to develop for four weeks (mean size approximately 150 mm^3^). Next, three doses of ROBO1 CAR T cells (1×10^6^, 3×10^6^, and 6×10^6^) were administered intravenously (n=4/group). Tumor growth was monitored by caliper measurements for up to 100 days or until humane endpoints were reached (Fig. S5A-F). The results showed a dose-dependent effect, with significant tumor regression and marked extended survival observed in mice with the two highest ROBO1 CAR T cell doses. The lowest dose was insufficient for tumor control. Based on these findings, we conducted an efficacy study (n=6/group) with 6×10^6^ ROBO1 CAR T cells, CD19 CAR T cells (negative control), or PBS (vehicle control) per mouse in both the 33T and 103T tumor models (Fig. 5A). Tumor growth was monitored by caliper measurements and bioluminescence imaging for over 50 days, and consistent with the dose-finding study, ROBO1 CAR T cells effectively suppressed tumor growth compared to the controls, resulting in extended survival and no tumor regrowth in both models (Fig. 5B-E, S6A-D). To assess CAR T cell persistence and expansion, we analyzed their presence in peripheral blood at days 7 and 14 post-injection using flow cytometry (Fig. 5F, S6E). At day 7, we observed similar numbers of CD19 and ROBO1 CAR T cells, confirming similar initial viability of the CAR T cells. However, by day 14, ROBO1 CAR T cells had significantly expanded, while CD19 CAR T cells had largely disappeared. This finding correlated with the infiltration of human CD45+ cells specifically in ROBO1-positive tumors of mice treated with ROBO1 CAR T cells, but not in those treated with CD19 CAR T cells or PBS (Fig. 5G, S6F).

**Figure 5.**
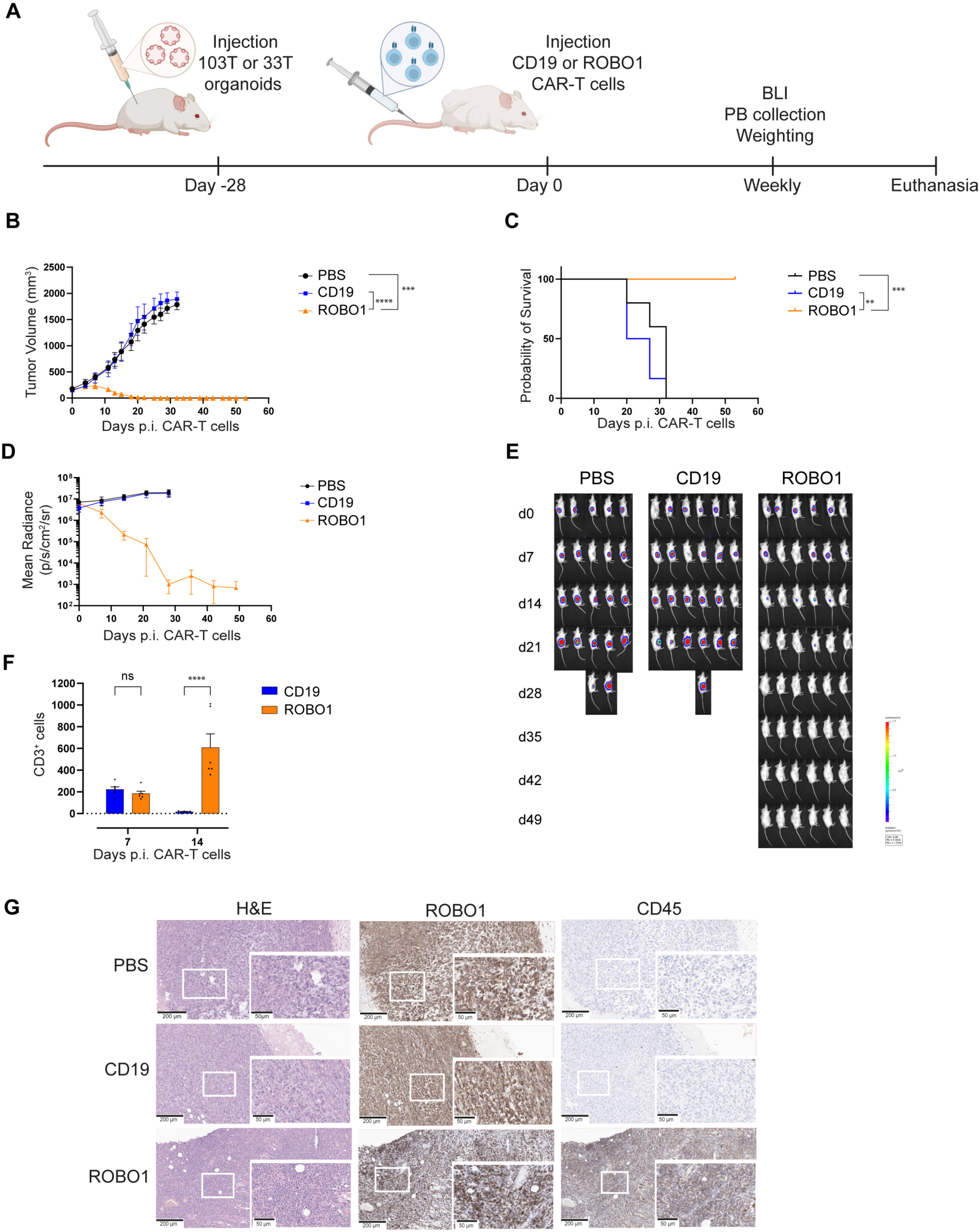
ROBO1-targeting CAR T cells induce potent and sustained cytotoxicity, eradicating eMRTs *in vivo*. (A) Schematics of the *in vivo* study. NSG mice were subcutaneously injected with luciferase-expressing eMRT organoids (103T or 33T). Acer 28 days of tumor engracment, mice received intravenous injection of either CD19 or ROBO1 CAR T cells (6 x 10^6^/mouse) at day 0. Tumor burden was monitored weekly using bioluminescence imaging (BLI). (B) Growth of 103T-engraced organoids acer CAR T cell treatment, shown as the mean tumor volume ± SEM measured by caliper (n=5 for PBS, n=6 for CD19 and ROBO1 CAR T cells). (C) Kaplan-Meier survival curves depicting overall survival of treated mice in (B). (D) Longitudinal tumor burden assessment from mice in (B) monitored by BLI. (E) Representative BLI images of tumor-bearing mice from (B), illustrating differences in tumor burden across treatment groups. (F) Flow cytometry analysis of peripheral blood (PB) from mice in (B), showing post-treatment immune cell counts. (G) Representative IHC images of tumor tissues, assessing human ROBO1 and human CD45 expression in treated mice. Statistical significance as in Fig. 2A.

In summary, this study highlights the therapeutic potential of ROBO1 as a CAR T cell target for MRT and other pediatric solid tumors, demonstrating its ability to support potent and sustained anti-tumor responses to CAR T cell therapy *in vitro* and *in vivo*.

## Discussion

Patients with MRTs continue to have a poor prognosis, with CAR T cell therapy offering promising new treatment possibilities. In this study, we aimed to address the scarcity of suitable CAR T cell targets with high translational potential for MRTs. An ideal CAR T cell target should be highly and uniformly expressed on the cell surface of tumor cells, with minimal or no expression in normal tissues. This ensures tumor specificity, maximizes efficacy, and minimize on-target, off-tumor toxicity (43). Although widely used in early preclinical drug development, classical cell lines fail to reflect the heterogeneity of the original patient tumors and therefore lack translational relevance. This issue is particularly important because the efficacy of CAR T cells is heavily influenced by antigen expression levels on the tumor cell surface, with the threshold for an efficient CAR T cell response varying depending on the antigen (28–30). To overcome these limitations and enhance the clinical translatability of our study to identify promising CAR T cell targets, we utilized patient-matched eMRT and normal kidney organoids developed in-house (12). These organoids closely mirror the transcriptional and genomic profiles of their parental tissues, providing a more accurate and physiologically relevant platform for target identification and preclinical evaluation. The growing recognition of the need for more representative models, such as patient-derived tumor organoids, in preclinical CAR T cell development, is evident in recent studies (25,49–52). Notably, a Phase I clinical trial involving six patients with recurrent glioblastoma demonstrated that tumor organoid–CAR T cell co-cultures, performed alongside treatment, could predict the patient’s response to dual-targeting CAR T cell therapy targeting EGFR and IL13Rα (53).

In this study, we performed cell surface proteomics on patient-matched MRT- and normal kidney tissue-derived organoids and, based on our selection criteria, identified four potential CAR T cell targets: ROBO1, TNFRSF10B, CSPG4, and MICB. Although MICB showed only weak overexpression in MRT organoids compared to normal kidney organoids, it was initially considered an interesting target for further investigation due to its known upregulation in response to cellular stress, such as DNA damage or malignant transformation, where it serves as a ‘kill me’ signal (54). Further validation, integrating (single-cell) transcriptomics and protein expression data, revealed that while CSPG4, MICB, and TNFRSF10B are upregulated in MRTs, their expression varies significantly both between patients and within individual tumors, suggesting that these may not be optimal CAR T cell targets.

In contrast, ROBO1 has emerged as a promising candidate for CAR T cell therapy due to its strong and uniform expression across eMRTs and ATRTs at both the RNA and protein levels. Although we have not quantified the surface density of ROBO1 molecules on MRT organoids, Chokshi *et al*. reported that ROBO1, overexpressed in glioblastoma, shows expression levels comparable to established CAR T cell targets such as B7-H3, CD70, IL13Rα2, and EGFR (46). Notably, ROBO1 has been shown to play context-depending roles in cancer biology. In some cancers, such as colorectal and hepatocellular cancer, it acts as an oncogene, promoting tumor growth and progression, while in others, including ovarian and cervical cancer, it functions as a tumor suppressor (34). Our study also found that ROBO1 is highly and uniformly expressed across several other pediatric solid tumors, including MRT, ATRT, GBM, HBL, NB, OS, RMS and WT, with more variable expression in EPD, EWS, and MB. Its broad expression in pediatric solid tumors underscores its potential for use in basket trials to evaluate ROBO1-targeting CAR T cell therapies, a critical consideration due to the rarity of pediatric solid tumors.

The heterogeneity of antigen expression observed in some of the pediatric cancers remains a significant challenge for CAR T cell therapy. Variable or low antigen expression can enable tumor cells to evade CAR T cell recognition, resulting in tumor escape, relapse and resistance to CAR T cells (55). To overcome this challenge, bispecific CAR T cell approach could be explored in a follow-up study. For instance, a ROBO1/B7-H3 bispecific CAR T cell design could be particularly effective, given that B7-H3 is also widely expressed in these tumor types (17). Various bispecific CAR formats, such as ‘tandem,’ ‘bispecific/bicistronic,’ and ‘logic gated’ designs, offer opportunities to optimize therapeutic outcomes (55,56).

A major challenge for CAR T cell therapy in solid tumors is on-target, off-tumor toxicity due to antigen expression on normal tissue (43). ROBO1, as an axon guidance receptor, plays a crucial role in neural and cortical development during embryogenesis, with ROBO1 knock-out mice exhibiting perinatal lethality (57). Although not classified as a bona fide oncofetal gene, publicly available data indicate that ROBO1 expression significantly decreases after birth and remains low in most adult tissues (44). Furthermore, several clinical studies are investigating the therapeutic potential of targeting ROBO1 in cancer. Phase I clinical trials (NCT03940820, NCT03941457, NCT03931720) are evaluating ROBO1-specific CAR-NK and CAR-NK/T therapies for solid tumors in adults, with safety results eagerly awaited. Furthermore, recent preclinical studies have shown ROBO1 overexpression in glioblastoma, medulloblastoma, and brain metastases (46). In these studies, ROBO1-targeted CAR T cells led to significant tumor reduction and improved survival in mouse models without causing toxicity or compromised neural integrity (46), consistent with our findings. Nevertheless, according to the Human Protein Atlas, ROBO1 is expressed in several non-malignant adult tissues, including the brain, gastrointestinal tract, and testis. However, the therapeutic window between ROBO1 expression in normal and cancerous tissue remains unclear. Evaluating the global expression of ROBO1, particularly across pediatric age groups, and characterizing the differential expression between normal and malignant tissues, remains challenging. While beyond the scope of the current study, PET imaging with target-specific probes could offer a comprehensive map of ROBO1 expression. Initial studies in mice of varying ages, with and without tumors, followed by human trials, could provide valuable insights. Furthermore, the development of immunoPET radiotracers for CAR T cell targets holds significant clinical relevance, as PET imaging is increasingly used for patient selection for immunotherapy (58). Such an approach could effectively assess the therapeutic potential of CAR T cells while also evaluating the risk of on-target, off-tumor toxicity prior to infusion, ultimately enhancing the safety and efficacy of ROBO1-targeted CAR T cell therapy. Once the therapeutic window of ROBO1 is better defined, the CAR design can be optimized. Strategies for refinement include tuning the CAR affinity to selectively target cancer cells, incorporating safety switches, using locoregional or local infusion methods, and other approaches aimed at minimizing off-tumor effects (43).

In summary, this study identifies ROBO1 as a highly promising target for CAR T cell therapy in MRTs and other pediatric tumors. Using patient-derived tumor organoids, which have proven effective in predicting patient responses to CAR T cell therapy, we show that ROBO1 is strongly and consistently expressed across various tumors. Furthermore, ROBO1-targeted CAR T cells promote robust and sustained anti-tumor efficacy, underscoring its potential for clinical translation.

## Material and Method

### Human tissues

Approval for use of human material was provided by the medical ethical committee of the Erasmus Medical Center (Rotterdam, the Netherlands) and the Princess Máxima Center for pediatric oncology (Utrecht, the Netherlands). Written informed consent was provided by all patients and/or parents/guardians. Approval for use of the subject’s tissue samples within the context of this study has been granted by the Máxima biobank and data access committee (https://research.prinsesmaximacentrum.nl/en/core-facilities/biobank); project numbers: PMCLAB2018.005, PMCLAB2018.006 and PMCLAB2024.0619.

### Cell lines

Cell lines were obtained from ATCC unless otherwise stated. HeLa cells were cultured in MEM (Gibco) supplemented with 10% heat-inactivated FBS (Sigma-Aldrich) and 1% Penicillin-Streptomycin (Gibco), A375, Huh7 and HEK293T cells were cultured in DMEM (Gibco) supplemented with 10% heat-inactivated FBS (Sigma-Aldrich) and 1% Penicillin-Streptomycin (Gibco), and Jurkat cells were cultured in RPMI-1640 (Gibco) supplemented with 10% heat-inactivated FBS (Sigma-Aldrich) and 1% Penicillin-Streptomycin (Gibco). Cell lines were regularly tested for mycoplasma contamination.

### Organoid cultures

#### MRT and normal kidney organoids

MRT and normal kidney organoids were previously established and characterized(12). The organoid lines were seeded in growth factor-reduced BME Type 2 (R&D Systems) and cultured in kidney organoid medium (Advanced DMEM/F12 (Gibco) containing 1X GlutaMAX (Thermo Fisher Scientific, #35050087), 10 mM HEPES (Thermo Fisher Scientific, #15630-056) and 1X Penicillin-Streptomycin (Merck Millipore, #516106) (AdDF+++), supplemented with 1.5% B-27 supplement (Gibco, #17504044), 10% R-spondin–conditioned medium, 50 ng/mLhEGF (PeproTech), 50 ng/mLhFGF-2 (PeproTech), 1.25 mM N-acetylcysteine (Sigma), 10 μM Rho-kinase inhibitor Y-27632 (Abmole) and 5 μM A83-01 (Tocris Bioscience). Medium was changed every 3-4 days and organoids were passaged using TrypLE Express (Gibco) with 10 μM Rho-kinase inhibitor once a week.

#### RMS organoids

RMS organoids were previously established and characterized(47). The organoid lines were cultured in the complete culture medium (BM1*). Base medium (BM) composed of advanced DMEM/F12 (Gibco) supplemented with 1X GlutaMAX (Thermo Fisher Scientific), 1X Penicillin-Streptomycin (Merck Millipore) and 2% B-27 supplement minus vitamin A (50×) (Gibco) was supplemented with 1% N2 (Gibco), 1% MEM nonessential amino acids (Gibco), 1 mM of Sodium pyruvate (Gibco), 20 ng/mL hEGF (Peprotech), 1.25 mM N-acetylcysteine (Sigma), 40 ng/mLhFGF-basic (Peprotech), 5 μM A83-01 (Tocris Bioscience), 0.5 U/mLHeparin (Sigma) and 10 μM Rho-kinase inhibitor Y-27632 (Abmole). To promote the attachment of cells 0.5% BME (R&D Systems) was added to the cool BM1*. Medium was changed every 3-4 days and cells were passaged using TrypLE Express (Gibco) once or twice a week depending on the growth rate.

#### NB organoids

NB organoids were previously established and characterized(48). The organoid lines were cultured in neuroblastoma medium, consisting of Dulbecco’s modified Eagle’s medium (DMEM) with low glucose and Glutamax™ (Gibco), supplemented with 20% Ham’s F-12 Nutrient Mix (Gibco), B-27 supplement minus (50X) (Gibco), N-2 supplement (100X) (Gibco), 1X Penicillin-Streptomycin (Merck Millipore), 20 ng/mL hEGF (PeproTech), 40 ng/mL hFGF-basic (PeproTech), 200 ng/mL hIGF-I (PeproTech), 10 ng/mL hPDGF-AA (PeproTech) and 10 ng/mL hPDGF-BB (PeproTech). Medium was changed every 3-4 days and cells were passaged by mechanical dissociation once or twice a week depending on the growth rate.

### CRISPR-Cas9-mediated knock-out of ROBO1 and luciferase transduction of organoid lines

Organoid knock-out clones were generated via CRISPR-Cas9 technology (59,60). Complementary single-strand oligonucleotides encoding the sgRNA sequence for ROBO1 knock-out (sgROBO1 sense oligo: CACCGTACACCCGTAAAAGTGACGC and sgROBO1 anti-sense oligo: AAAC GCGTCACTTTTACGGGTGTAC) were annealed and cloned into the lentiCRISPRv2 puro vector (Addgene, 52961) via BsmBI-v2 (NEB, R0739) restriction and T4 ligation (NEB, M0201) based on previously described protocols of the Zhang lab. Luciferase transduction was performed as previously described(61) using the pLKO.1-Ubc-luciferase-blast plasmid. Lentivirus vectors were produced in HEK239T cells. The cells were transiently transfected with pHDM-G, pRC/CMV-rev1b, pHDM-Tat1b, pHDM-Hgpm2 and lentiCRISPRv2-sgROBO1 using 25 kDa polyethylenimine (PEI) (Tebubio, 23966-1). Medium was replaced after 24 h and virus supernatant was harvested after 72 h. The supernatant was filtered using a 0.45 μm filter, centrifuged at 20,000 x g for 1.5 h at 4°C. The pellet of one 15 cm culture dish was resuspended in 500 μl growth media, snap frozen and stored at -80 °C. Transduction of organoids was performed after dissociation into single cells with concentrated lentiviral supernatant in the presence of 1:1,000 polybrene (10 mg/mL, SantaCruz). After spinoculation at 32°C at 600 x g for one hour, cells were incubated at 37°C for 4 hours, subsequently washed and reseeded in BME droplets. 2 days after transfection, 1 μg/mL puromycin (Invivogen) or 10 μg/mL blasticidin (Invivogen) selection was started and carried out for up to 2 weeks. Finally, to generate ROBO1 knock-out clones, single cells were sorted into a 96-well u-bottom plate, cells allowed to grow out and the knock-out was confirmed on protein level by flow cytometry and DNA level by sequencing.

### Cell surface biotinylation of organoid lines

For selective enrichment of cell surface proteins expressed on MRT and normal kidney organoid lines, Pierce Cell Surface Protein Biotinylation and Isolation Kit (Thermo Fisher Scientific, #A44390) was used. MRT organoid lines 60T and 103T and normal kidney organoid lines 60H and 103H were expanded in kidney organoid medium to ∼20×10^6^ cells per line (1 6-well MRT organoids yields ∼2.5×10^6^ cells, and 1 6-well normal kidney organoids ∼1.0×10^6^ cells). Since these organoids are neither adherent nor in suspension, the protocol of the kit was modified. Organoids were harvested from the culture plates (three 6-wells per tube) in DMEM (Gibco) without disrupting the organoid structure and spun down for 5 min at 300 x g and 4°C with half the deceleration speed. To remove the remaining BME, the pellet was resuspended in 1 mL of collected supernatant and 9 mL of supernatant was added. Organoids were spun down for 5 min at 300 x g and 4°C with half the deceleration speed. This step was repeated. Cell pellets were washed two times with ice-cold PBS using low-binding tips. Organoids were spun down for 3 min at 300 x g and 4°C with half the deceleration speed and the supernatant was discarded.

For cell surface protein biotinylation, organoids were resuspended in 2 mL/tube of freshly prepared EZ-Link Sulfo-NHS-SS-Biotin solution in PBS (6 mg/10×10^6^ cells), incubated for 15 min on ice and gently mixed every 3 min. Organoids were then spun down for 3 min at 300 x g and 4°C with half the deceleration speed and the supernatant was discarded. Cells were washed twice with 5 mL of ice-cold TBS. Cells were spun down for 3 min at 200 x g and 4°C with half the deceleration speed and the supernatant was discarded. Cells were resuspended in 2 mL/tube ice-cold TBS and the tubes were combined. Cells were spun down for 3 min at 200 x g and 4°C with half the deceleration speed and the supernatant was discarded. Cells were resuspended in 4 mL of ice-cold TBS, of which 150 μL was used for immunofluorescence staining. Cells were spun down for 3 min at 200 x g and 4°C with half the deceleration speed and the supernatant was discarded.

To solubilize proteins, organoids were first lysed by gently vortexing in 2 mL hypotonic lysis buffer (10 mM Tris buffer (pH 7.5) with 1 mM PMSF (Sigma-Aldrich, #52332, dissolved in isopropanol) supplemented with 1X EDTA-free protease inhibitor cocktail (Thermo Fisher Scientific, #A32955)) every 2 min for 10 min at 4°C, after which they were transferred to low binding Eppendorf tubes. Subsequently, the lysate was cleared by centrifugation for 15 min at 16,000 × g and 4°C. Supernatant was collected as cytosolic fraction 1 for Dot blot analysis. The pellets were resuspended in 2 mL hypotonic lysis buffer and gently vortexed every 2 min for 10 min at 4°C. Lysates were spun down for 15 min at 16,000 × g and 4°C. Supernatant was collected as cytosolic fraction 2 for Dot blot analysis. The pellets were resuspended in 600 μL lysis buffer and lysis was repeated by vortexing every 10 min for 30 min at 4°C. The lysate was cleared by centrifugation for 20 min at 16,000 × g and 4°C.

For isolation of biotinylated proteins, the suspension with the labeled cell surface proteins was collected, of which 100 μL was used to determine total protein amount with the BCA protein assay (Thermo Fisher Scientific, #23225) and input sample for Dot blot analysis. Based on the protein concentration, the required volume of NeutrAvidin agarose was determined for each sample and added to the spin columns provided by the kit. The columns were spun 1 min at 1,000 x g and the flow-through was discarded. To prepare the resin for binding, the resin was washed three times with wash buffer (volume was adjusted for each sample based on resin volume and protein sample), and spun for 1 min at 1,000 x g and the flow-through was discarded. 500 μL biotinylated cell surface protein fractions were then added to the prepared columns. Samples were incubated for 30 min at RT with end-over-end mixing using a rotator. The NeutrAvidin columns were placed into Eppendorf tubes and spun for 1 min at 1,000 x g and the flow-through was collected as flow through sample for dot blot analysis. The columns were washed three times with wash buffer A (wash buffer containing 1 mM PMSF and supplemented with 1X EDTA-free protease inhibitor cocktail) and two times with wash buffer B (TBS with 1 mM PMSF supplemented with 1X EDTA-free protease inhibitor cocktail). For each wash step, one volume (dependent on the amount of protein loaded) was added to the column after which the column was spun for 1 min at 1,000 x g and the flow-through were collected as Wash buffer A 1, 2 and 3 and Wash buffer B 1 and 2 for dot blot analysis.

For immunofluorescence staining, biotinylated organoids were washed with cold PBS and spun down for 3 min at 300 x g and 4 °C. The organoids were gently resuspended in 1 mL of 2% PFA and incubated for 45 min at 4 °C. Organoids were washed with PBST and spun down for 5 min at 250 x g and 4 °C. The organoids were blocked for 15 min at 4 °C with Organoid Washing buffer (OWB; PBS containing 0.1% Triton X-100 and 0.2% BSA) and transferred to a low-adherence 24-well plate. After organoid settling at the bottom of the plate, half of the OWB was removed and Streptavidin-AF647 (Final concentration 1:500) was added and incubated overnight at 4 °C with mild shaking (60 rpm). OWB (1 mL/well) was added to the organoids after organoid settling at the bottom of the plate, half of the OWB was removed. Organoids were washed three times with 1 mL/well of OWB for 30 min at 4 °C with mild shaking. DAPI (final concentration 300 nM) was added and incubated for 10 min at RT. Organoids were washed with 1 ml/well PBS for 3 min followed by three times washing with 1 mL/well PBS for 5 min with mild shaking. Organoids were imaged using an SP8 Confocal microscope (Leica) with a 10x/0.40 dry objective and analysis was performed with LAS X 3.5.6.21594 software (Leica).

For dot blot analysis, the volume that needs to be spotted on the nitrocellulose membrane for each sample (1 μg of protein for biotin staining and 5 μg for CD29 or ß-actin staining) was determined by the BCA protein analysis of the input sample as described above. After spotting, the membrane was air-dried. The membrane was blocked with Blocking buffer (TBS supplemented with 5% BSA and 0.05% Tween20) for 1 h at RT. After blocking the membrane was incubated with primary antibody, CD29 (1:1000, Invitrogen, 14-0299-82) or ß -actin (1:200, Santa Cruz, sc-47778), diluted in Antibody buffer (TBS supplemented with 0.1% BSA and 0.05% Tween20) for 1 h at RT. After incubation, membranes were washed three times with TBST (TBS supplemented with 0.05% Tween20). Membranes were incubated with Rabbit anti-mouse-HRP (1:2000 or 1:1000, respectively, Invitrogen, 616520) or ExtrAvidin-HRP (1:4000, Sigma-Aldrich, E2886-0.2mL) diluted in Antibody buffer for 1 h at RT. After incubation, membranes were washed three times with TBST and once with TBS. The immunoblots were visualized using enhanced chemiluminescence detection (ECL Plus Substrate, Thermo Scientific, 32132).

### Liquid chromatography with tandem mass spectrometry (LC-MS/MS)

Cell surface biotinylated proteins captured by NeutrAvidin pulldown were on-column reduced by 50 mM Dithiothreitol (DTT) for 1 h at 20°C and then alkylated with 100 mM iodoacetamide (IAA) for 0.5 h at 20°C in the dark. Proteins were then digested sequentially at 37°C by 0.5 µg of Lys C and trypsin, for 4 h and 12 h, respectively, in buffer containing 2 M Urea and 50 mM ammonium bicarbonate. Digested peptides were acidified to pH<3.0 for further double purification by strong cation exchange using the Supel™-Select SCX SPE Tubes (Supelco) and C18 (Empore solid phase extraction discs) STAGE tips. Eluted peptides were dried by vacuum centrifugation. Peptides were reconstituted in 2 % formic acid and triplicate injections of 25 % each were analyzed on an Orbitrap Exploris 480 mass spectrometer (Thermo Scientific), coupled to an UltiMate 3000 UHPLC system (Thermo Scientific). Solvents used were 0.1 % formic acid in water (Solvent A) and 0.1 % formic acid in 80 % acetonitrile, 20 % water (Solvent B). Peptides were first trapped on an µ-precolumn (C18 PepMap100, 5 µm, 100 Å, 5 mm × 300 µm; Thermo Scientific) in 9 % Solvent B, and then separated on an analytical column (120 EC-C18, 2.7 µm, 50 cm × 75 µm; Agilent Poroshell) using a 95 min linear gradient of solvent B per fraction. The resolving gradients for fractions 1 to 5 were 11-20 %, 12-28 %, 15-34 %, 18-35 % and 23-40 % Solvent B, respectively. Eluting peptides were online-injected into the mass spectrometer for data-dependent acquisition. The spray voltage was set to 2.1 kV, the temperature of the ion transfer tube was set to 275°C and an RF lens voltage of 40 %. MS scans were acquired at a resolution of 60,000 within the m/z range of 375-1600, accumulating to ‘Standard’ pre-set automated gain control (AGC) target. Multiply charged precursor ions starting from m/z 120 were selected for further fragmentation. Higher energy collisional dissociation (HCD) was performed with 28 % normalized collision energy (NCE), at a resolution of 30,000, and with dynamic exclusion of 16 s and 1.4 m/z isolation window.

### Quantitative proteomics data analysis

MS data was acquired with Thermo Scientific Xcalibur version 4.4.16.14, and raw files were processed using MaxQuant software version 1.6.15.0 with the integrated Andromeda search engine. Data were searched against the human UniProt database (downloaded in March 2020, containing 188,357 entries) including common contaminants. For all files standard parameter settings were used with enabled the label-free quantification (LFQ) algorithm. Cysteine carbamidomethylation was included as a fixed modification. Protein N-terminal acetylation and methionine oxidation were allowed as variable modifications. Trypsin/P was set as the digestion enzyme (cleaves after lysine and arginine also if a proline follows), and up to two missed cleavages were tolerated. The match-between-run feature was enabled for identification. A false discovery rate (FDR) of 1 % was used for peptide and protein identification. Quantitative data were analyzed using Perseus software version 1.6.10.0. LFQ intensities of proteins were log2-transformed. Proteins quantified in two out of three replicates in one of experimental conditions were retained for further analysis, after imputation based on normal distribution. From this list of proteins, we applied a stringent filtering workflow comprising of information from UniProt such as Gene Ontology (GO) terms: plasma membrane (GO:0005886), apical plasma membrane (GO:0016324), basolateral plasma membrane (GO:0016323), lateral plasma membrane (GO:0016328), cell surface (GO:0009986) and external side of plasma membrane (GO:0009897), subcellular localization terms (cell membrane, apical cell membrane, basolateral cell membrane, transmembrane, intramembrane and GPI-anchor), the presence of signal peptides as well as cell surface protein information from the SURFY database to discern potential cell surface targets. To identify differentially expressed cell surface proteins between experimental conditions, Student’s T tests were performed. FDR-corrected p-values (q-values) were calculated from 250 randomizations and considered significant if they were 0.05 or less.

### Evaluation of antigen expression in available bulk RNA sequencing data of eMRT and ATRT organoids and combined single-nuclei RNA and ATAC sequencing datasets

The processed bulk RNA sequencing data of ATRTs and eMRTs (tissues and organoids) from the EGA database (EGAS00001006866: ATRT tissues n=7, ATRT organoids n=9; EGAS00001003853: MRT tissues n=10, MRT organoids n=7, normal kidney organoids n=5) were used in this study (12,14). Raw counts were converted to counts per million (CPM) log2 transformed. Normalization was performed using the trimmed Mean of M-values (TMM) method. The processed sequencing data of combined single-nuclei RNA and ATAC data (9) (eMRT tissues n=7) from the GEO under accession code <u>GSE218385</u> were used to map target expression. UMAP plots were generated via joined clustering of both datasets (PCA 1:25, LSI 2:30).

### Flow cytometry of patient-derived organoids, CAR T cells and commercial cell lines

For evaluating antigen expression on MRT, RMS and normal kidney organoids, the organoids were harvested and dissociated into a single-cell suspension using TrypLE (Gibco, 12605010) supplemented with Rho-kinase inhibitor Y-27632. For NB organoid lines, the organoids were either dissociated mechanically or using Accutase (Thermo Fisher), depending on the cell line. All cells were counted, washed twice with FACS buffer (PBS containing 2 % FBS and 1 mM EDTA) and 1.5 x 10^5^ cells/well were transferred to U-bottom-shaped 96-well plate. For evaluating the binding of anti-ROBO1 scFv-Fc to cell lines, HeLa, Huh7 and A375 cells were dissociated from culture flasks into a single-cell suspension with 1 mM Sodium Citrate and 1 mM EDTA in PBS, washed twice with FACS buffer and transferred to V-bottom-shaped 96-well plate (Sigma Aldrich, M9561-100EA) at 1 x 10^5^ cells/well. Jurkat cells were cultured in suspension and were collected and washed twice with FC buffer before they were transferred to V-bottom 96-well at 1 x 10^5^ cells/well.

All subsequent steps were performed on ice, all scFv or antibody incubation steps were done for 1 h with shaking and all washing steps were performed twice with FACS buffer and centrifugation of the plate at 300 x g for 5 min at 4°C. Cells were incubated with ROBO1-targeting scFv-Fc-LC-biotin (production described below), specific primary antibodies (TNFRSF10B (1:500, Sanbio B.V., Ab00740-10.0), ROBO1 (1:200, R&D, MAB71181), MICB (1:2000, R&D, MAB1599) and CSPG4 (1:2000, R&D, MAB2585)) or corresponding isotype control (Mouse IgG1,k (Invitrogen, 14-4714-81), Mouse IgG2b (Invitrogen, 02-6300) or Human IgG1,l (Arigo Biolaboratories, ARG20768)) and detected with Alexa Fluor 647-labelled streptavidin (1:500, Invitrogen, S32357) or secondary anti-mouse or anti-human antibodies (1:500, Invitrogen, A21235 and 1:1000, Invitrogen, A21445, respectively). Live/Dead staining was done with DAPI (1:500, Invitrogen, D1306).

To determine activation of CAR T cells, CAR T cells and MRT organoids were co-cultured in a 1:10 effector to target ratio for 20 hours. Cells were washed with FACS buffer twice and incubated with hCD45-FITC (1:100, Thermo Fisher) and CD137-APC (1:25, BD Bioscience) for 30 min at 4°C. After washing the cells with FACS buffer twice, DAPI (1:500) was added and cells were measured at the flow cytometer.

Cells were analyzed using the Beckman Cytoflex S or LX flow cytometer and the results were analyzed using FlowJo^TM^ v10 Software (BD Life Sciences). A minimum of 5000 live cells were used for analysis after Live/Dead gating.

### Western blot

MRT and normal kidney organoids were harvested, counted and lysed with RIPA lysis buffer (150 mM NaCl, 1 % Triton X-100, 0.5 % Sodium deoxycholate, 0.1 % SDS, 50 mM Tris-HCl, pH 8.0) containing protease inhibitors (1 mM PMSF and protease inhibitor mini tablets, EDTA-free (Thermo Scientific)) at 4°C for 30 min. Samples were sonicated and total protein was collected following a 30 min centrifugation at 16,000 x g at 4°C. BCA protein assay (Thermo Scientific) was used to determine protein concentration. A total of 20 μg of protein was electrophoresed in a 7.5 % SDS-PAGE gel. The proteins were transferred to 0.45 μM polyvinylidene difluoride (PVDF) membranes (Immobilon-FL, Merck) and blocked with 5 % skim milk in Tris-buffered saline (TBS) containing 0.1 % Tween 20. The membranes were incubated at 4°C overnight with specific primary antibodies (TNFRSF10B (1:7000, ProSci, 2019), ROBO1 (1:8000, Proteintech, 20219-1-AP), MICB (1:3790, R&D, AF1599), CSPG4 (1:4000, Cell signaling, 43916T) or GAPDH (1:150,000, Proteintech, 60004-1-Ig)), followed by 1 h incubation with horseradish-peroxidase-conjugated secondary antibodies (anti-Rabbit-HRP (1:50,000, Abcam, AB205718), anti-Goat-HRP (1:30,000, Invitrogen, 31400) or anti-Mouse-HRP (1:10,000, Invitrogen, 31430). The immunoblots were visualized using enhanced chemiluminescence detection (ECL Plus Substrate, Thermo Scientific).

### Bioinformatic analysis M&M dataset

For gene expression analyses performed for Fig. 4A-B and Fig. S4A-B, RNA-seq data was obtained from the publicly available repository hosted on Zenodo (62) related to the publication of the M&M classifier (42). Data was subsetted to only include eMRT, ATRT, RMS, NB, EWS, WT, MB, OS, GBM, HBL or EPD samples. Counts per million (CPM) where log10 (CPM+1) transformed to enhance visualization as recommended in previous transcriptomic studies (63). Data processing, normalization and visualization were conducted in R (version 4.4.2) (64) using the ggplot package (65).

### Immunohistochemistry (IHC)

To demonstrate the overexpression of CSPG4, MICB and ROBO1 in FFPE tissue of rhabdoid tumors, a total of 22 different cases were included for which the diagnosis of rhabdoid tumor was confirmed. These rhabdoid tumors could be subdivided into ATRT (n=5), MRT (n=15) and metastatic rhabdoid tumors (n=2). The IHC analysis includes FFPE tissues from 17 resection specimens and 5 biopsies from rhabdoid tumor patients. IHC was performed on normal tissues (pediatric and adult) commonly affected by MRTs: the kidney (n=14), liver (n=10) and brain (n=4). Several differential diagnoses have been investigated in order to examine the specificity of the biomarkers for rhabdoid tumors. To expand the evaluation of target expression to other tumor entities, we also included neuroblastoma (n=3), renal cell carcinoma (n=3), Wilms tumor (n=7), of which 4 were metastatic Wilms tumors, medulloblastoma (n=2), rhabdomyosarcoma (n=4), and Ewing sarcoma (n=2). For the MRT PDX tumor samples one tumor per treatment condition and organoid model was assessed.

Three micrometer thick sections of FFPE blocks were cut, mounted on precoated slides, and dried for at least 30 min at 65°C. For biomarkers CSPG4, MICB and ROBO1, as well as hCD45 deparaffinization and IHC staining was performed with an automated BOND-RX system (Leica Microsystems). Antigen retrieval was performed by boiling the sections in Tris/EDTA (BOND Epitope Retrieval Solution 2, pH9; Leica Biosystems) for 15 min for ROBO1 and 20 min for CSPG4 and MICB. For human CD45, antigen retrieval was performed by boiling the sections for 20 min in citrate-based buffer (BONE Epitope Retrieval Solution 1, pH 6; Leica Biosystems). The following primary antibodies were used: ROBO1 (clone 10E2-R, Invitrogen Antibodies, 1:400, retrieval buffer Tris/EDTA pH9); CSPG4 (clone EPR9195, Abcam, 1:400, retrieval buffer Tris/EDTA pH9); MICB (clone EPR2203, Abcam, 1:600, retrieval buffer Tris/EDTA pH9); human CD45 (clone X16/99, Leica, ready to use, retrieval buffer citrate pH6). The primary antibodies were diluted with BondTM Primary Antibody Diluent (Leica). The sections were incubated at room temperature for 15 min for hCD45, 25 min for MICB and 30 min for CSPG4 and ROBO1. The sections were then incubated for 8 min with a post-primary rabbit anti-mouse linker followed by incubation for 8 min with anti-rabbit horseradish peroxidase-labeled polymer. Endogenous peroxidase was blocked with 0.3 % hydrogen peroxide in deionized water for 5 min. After incubation for 10 min with diaminobenzidine (DAB) and again 5 min with DAB enhancer, all sections were counterstained with hematoxylin (BOND Polymer Refine Detection Kit; Leica Biosystems) for 5 min, dehydrated, cleared and mounted. A positive and negative control tissue was included in each section. Under the Leica DM 2500 light microscope (Leica Co), the stained sections were scored according to the four-point scale based on the IHC intensity.

IHC scoring was performed blinded by two people, the diagnostics department team leader and an analyst specialized in IHC staining. A semi-quantitative scoring system was used for the assessment based on the staining intensity with a minimum staining of >10 % of the tumor cells. Staining intensity: 0 = no staining; 1 = weak; 2 = moderate; 3 = strong.

### Expression and purification of anti-ROBO1 scFv-Fc

ROBO1 scFv sequence, from the patent US 2022/0267731 A1, was cloned into a pcDNA3.1 vector with a CH2-CH3 (Fc) domain of human IgG1 on the C-terminus. The expression was performed in Expi293F cells by transient transfection of the plasmid using the ExpiFectamine 293 transfection kit (ThermoFisher, A14635) according to the manufacturer’s protocol. After 4 days the cell cultures were centrifuged and the medium containing recombinant scFv-Fc was harvested. The scFv-Fc was captured from the cleared supernatant using CaptivA Protein A Affinity Resin (Repligen, CA-PRI-0100) and eluted with IgG elution buffer (Thermo Scientific, 21009) containing 500 mM NaCl and neutralized with 100 mM Tris pH 8.0 to reach eventually pH 7.0. Next, size exclusion chromatography (SEC) with Superdex^TM^ 200 Increase 10/300 CL column (Cytivia) on the ÄKTA pure^TM^ chromatography system (Cytivia) with HEPES-buffered saline (HBS) supplemented with 500 mM NaCl was performed. Fractions containing the scFv-Fc were pooled and concentrated using Amicon centrifugal filters (Millipore, UFC8030) and sterilized by filtration over a 0.22 μm centrifugal filter (Merck, UFC40GV0S). Protein concentrations were determined via NanoDrop A_280_ readings using the respective protein molecular weight and extinction coefficients. For subsequent experiments, the scFv-Fc was chemically biotinylated. For this, sulfo-NHS-LC-Biotin (Thermo Scientific, A39257) was dissolved in H_2_O and incubated in a 20:1 Dye : Protein molar ratio for 1 h at RT. Unbound dye was removed with a Zeba spin 7 kDa desalting column (Thermo Scientific, 89882) according to the manufacturer’s protocol.

### Enzyme-linked immuno sorbent assay (ELISA)

Maxisorp plates were coated with 5 μg/well recombinant hROBO1 (R&D, 8975-RB) in PBS overnight at 4°C. The next day, plates were washed with PBST and blocked with 2% Skim milk in PBST (blocking buffer) for 1 h at RT with shaking. ROBO1-targeting scFv-Fc-LC-biotin was diluted in blocking buffer, incubated, and detected with HRP-conjugated ExtrAvidin (Sigma-Aldrich, E2886). All the incubations were done for 1 h at RT while shaking and between all incubation steps the wells were washed three times with PBST. The binding was determined with TMB substrate reagent (BD, 555214) and the reaction was stopped with 1 N HCl. Optical density was measured at 450 nm with a SPECTROstar Nano microplate reader (BMG Labtech). The data was analyzed using Graphad Prism v9 Software.

### ROBO1 and CD19 CAR T cell production

Engineered CAR T cells were produced according to a previously published protocol(66). The CAR19 construct was established as described previously (67). The utilized ROBO1 scFv was derived from patent US 2022/0267731 A1 and cloned into the CAR19 backbone. Blood was acquired via the UMC Utrecht Minidonordienst, the collection protocol was approved by the ethical committee of the UMC Utrecht. PBMCs were isolated from peripheral blood by Ficoll-Paque PLUS separation (GE-healthcare, #17-1440-03). According to manufacturer’s protocol, CD3+T cells were isolated using αCD3-based magnetic bead separation (Miltenyi Biotec, #130-050-101). T cells were expanded for 3 days using αCD3/CD28 dynabeads (Gibco, ThermoFisher) in a ratio of 1:2.5 (bead to cell) in TCM: RPMI supplemented with 5% human serum, 1% penstrep supplemented with 50 U/mL IL-2 (provided by Pharmacy UMC), 5 ng/mL IL-7 (Miltenyi Biotec, #130-095-563) and 5 ng/mL IL-15 (Miltenyi Biotec, #130-095-765). Then, T cells were either left untransduced or transduced with the ROBO1-or CD19 CAR-construct containing virus. For this, 10 μl of virus per condition was added to 0.5 x 10^6^ T cells in 200 μl of Stem-MAX medium containing Lentiboost A+B (Sirlion Biotech, 1:100) in a 96-well U-bottom plate and left in the incubator at 37°C for 24 hours. For T cells used in the *in vivo* experiment, cells were transduced at a multiplicity of infection of 10 with the titer determined using Jurkat cells, as described previously (68). After transduction T cells were seeded in T cell medium (TCM) containing RPMI (Gibco, #72400047) + 10% FCS (Vidavi, #134419) + 1% Pen/Strep (Gibco, #15140122) with 50 U/mL IL-2 (Miltenyi Biotec, #130-097-744), 5 ng/mL IL-15 (Peprotech, #200-15) and 5 ng/mL IL-7 (Peprotech, #200-07). Then the transduction efficiency was determined using Flow Cytometry. For this, 50000 of the transduced cells were stained with FITC-conjugated anti-murine-IgG (Sigma Aldrich, #F9137; 1:100) for 20 min on a shaker at 4°C. To isolate transduced from non-transduced T cells, the cells were FACS-sorted on SONY (SH800S cell sorter) based on their ROBO1 or CD19 construct expression. For this, cells were stained with CD3-AF-700 (1:200, Biolegend, #300323), and ROBO1 conjugated with FITC protein (1:100, Acrobiosystems, #RB1-HF2H7) or CD19-FITC protein (1:100, Acrobiosystems, #FM3-FY45). The sorted cells were collected in human serum and expanded based on a previously described protocol (69). In short, 1 x 10^6^ T cells were cultured in RPMI-Glutamax (Gibco) supplemented with 5% pooled human serum (≥3 healthy donors), Beta-2-mercaptoethanol, and 1% Pen/Strep (Gibco) on a cell mixture of 2.5 x 10^6^ cryopreserved irradiated (8000 cGy) pooled LCLs derived from three healthy donors (Coriell Institute for Medical research) and 12.5 x 10^6^ cryopreserved irradiated (3500 cGy) pooled PBMCs from ≥3 healthy donors in the presence of IL-2 50 U/mL (Miltenyi Biotec, #130-097-744) and 1 μg/mL PHA-L (Thermofisher Scientific). After 4 days, PHA-L was washed out and cultures were replenished with fresh culture media and 50 U/mL IL-2 was added. After 7 and 10 days of culture, a half media change was performed and 50 U/mL IL-2 was added. After T cell expansion quality was controlled using Flow Cytometry to identify proportions of the CD4+ and CD8+ T cells as well as the expression of the CAR using CD3-APC (1:100, Biolegend, #300412), CD4 Violet blue (1:50, Miltenyi biotec, #130-113-219), CD8-V500(1:50, BD, #561617) and ROBO1-FITC protein or CD19-FITC protein. The analysis was performed on Cytoflex S and FlowJo as described above.

### CAR T cell - tumor organoid co-culture

To determine tumor cell specific killing, all organoid lines were transduced with the pLKO.1-UbC-luciferase-blast by lentiviral transduction as previously described(60,61,70,71). Two days after transduction, transduced cells were selected by the addition of 10 µg/mL blasticidin (Invivogen).

For MRT, organoids were cultured in kidney organoid medium in BME droplets for four days prior to the start of the experiment. On the day of the co-culture, MRT organoids were carefully collected, washed with cold medium to remove BME, and a fraction was dissociated into a single cell suspension. Single tumor cells were counted to determine the number of cells present in the MRT organoid suspension. MRT organoids were then seeded out with an equivalent of 7500 single tumor cells per well in a 96-well plate in 50 µL of co-culture medium containing 10 % FBS and a 1:1 ratio of kidney organoid medium and RPMI with 1X GlutaMAX and 1X P/S. Technical replicates were performed in quadruplicates. ROBO1 and CD19-directed CAR T cells were defrosted three days in advance and seeded out in RPMI containing 1X GlutaMAX (Gibco), 1X P/S and 10% FBS supplemented with 50 IU/mL IL-2, 5 ng/mL IL-7 and 5 ng/mL IL-15 in a density of 1 x 10^6^ cells/mL. On the day of the co-culture, CAR T cells were washed with RPMI and counted. For the 1:10 ratio 750 cells, for the 1:25 ratio 300 and for the 1:50 ratio 150 live CAR T cells were added in 50 µL of co-culture medium per well to the 96-well plate. To determine tumor-specific killing, a luciferase assay was performed using the luciferase assay system (Promega). Medium was removed after 24 h and stored at -80°C for cytokine secretion analysis.

For the detection of the luciferase signal, cells were first washed with PBS, then 20 µL of 1X passive lysis buffer (Promega) were added and the plate was incubated at room temperature for 15 min while shaking. In a LUMITRAC white polystyrene 96-well plate (Greiner) 50 µL of the luciferase reagent were added to 20 µL of the lysed cells and the plate was measured at the FluoSTAR Omega Microplate reader immediately. Luciferase signal was determined on day 1, day 3 and day 5.

For RMS, tumor cells were re-seeded four days in advance in BM1* medium. On the day of co-culture, cells were washed with PBS, detached with TrypLE and counted. Depending on the proliferation speed of the organoid lines, 5000 cells of line RMS444 were seeded per well and 7500 cells of RMS013 and RMS000HQC were seeded out in 50 µL co-culture medium containing 0.25 % BME, 10 % FBS and a 1:1 ratio of BM1* and RPMI with 1X GlutaMAX and 1X P/S. CAR T cells were added a described above and cell numbers adjusted to the numbers of seeded RMS tumor cells. Tumor cell killing was determined as described above on day 1 and day 3.

For NB, organoids were re-seeded four days in advance in NB medium. On the day of co-culture cells were mechanically detached, washed and as small fraction made single cell to count. 30,000 tumor cells were seeded per well in 50 µL of co-culture medium containing 10 % FBS and a 1:1 ratio of NB medium and RPMI with 1X GlutaMAX and 1X P/S. CAR T cells were added a described above and cell numbers adjusted to the numbers of seeded NB tumor cells. Tumor cell killing was determined as described above on day 1 and day 3.

### Live-cell imaging

Live-cell imaging was performed as described previously with minor modifications(50). In brief, to visualize CAR T cells, cells were washed with PBS and stained with eBioscience Cell Proliferation Dye eFluor 450 (Thermo Fisher) in PBS in a 1:1000 dilution for 10 min at 37°C. RPMI with 10% FBS was added, and cells were further incubated on ice for 5 min. Then cells were washed, counted and seeded out in black, glass-bottom 96-well plates (Greiner) together with MRT organoids in a E:T ratio of 1:10 ratio (7,500 CAR T cells: 37,500 MRT cells) in 200 μL co-culture medium containing 5% BME and NucRed Dead 647 (2 drops/mL, Thermo Fisher). Cells were incubated in the incubation chamber of the Leica TCS SP8 confocal microscope (5% CO2, 37°C) and imaged every 30 min with a HC PL APO CS2 10x/0.4 dry objective for 72 h. The following settings were used: resonant scanner at 8000 Hz, bidirectional scanning, line averaging of 8, frame accu of 2, 3 µm Z-stack step size, Z-stacks of 110 µm in total and a resolution of 512 x 512 resulting in a Voxtel size of 1.82 µm x 1.82 µm x 2.409 µm. Analysis was performed using Imaris (Oxford Instruments) v.9. and median filter 3 x 3 x 3 was applied to the 633 nm laser signal.

### Multiplexed immunoassay

Supernatants were collected after 24 h co-culturing of CAR T cells and MRT organoids. Assay was performed according to the manufacturer’s protocol (LEGENDplex Human CD8/NK Panel V02, Cat. No. 741187). All the incubation steps were performed at room temperature on a shaker with a speed of 800rpm. Briefly, assay buffer, mixed beads and diluted standards or samples were incubated for 2 h. The plate was centrifuged at 250 x g for 5 min and the supernatant was removed by quickly inverting and flicking the plate. After washing with provided wash buffer, detection antibodies were added and incubated for 1 h. Subsequently, SA-PE was added and incubated for 30 min. The plate was again centrifuged at 250 x g for 5 min and supernatant was removed. The plate was washed again. In the end, wash buffer was added to resuspend the beads and signals were detected using a flow cytometer.

### *In vivo* mouse studies

All animal experiments were performed after approval and in accordance with the guidelines of the Ethical Committee for Experimental Animals at the Faculty of Health and Medicine and Health Sciences of Ghent University (ECD 23-06, Ghent, Belgium). Six- to ten-week-old male or female in-house bred NOD.Cg-PrkdcscidIL-2rgtm1Wjl/SzJ (NSG) mice were used in all *in vivo* experiments. 33T and 103T organoid cells, 2.5 x 10^5^ and 5 x 10^5^ respectively, were subcutaneously injected with a 30g needle (BD Biosciences) in 100 µl of volume. Before injection, the cells were resuspended in 100 µl of a 1:1 mixture of kidney organoid medium and ice-cold BME. Tumors were allowed to form for four weeks after which 1, 3 or 6 x 10^6^ (numbers based on transduced T cells) thawed CD19- or ROBO1-specific CAR T cells were injected into the tail vein. All injected cells were resuspended in PBS and a maximum volume of 200 µl was administered for tail vein injection. Tumor burden was defined by caliper measurements of tumor dimensions in mm and calculated using the formula length² x width/2 and bio-luminescence measurements. Bioluminescence was assessed after intraperitoneal injection of 150 mg/kg bodyweight D-luciferin (Perkin Elmer) using an IVIS Lumina III *in vivo* imaging system (Perkin Elmer). Animals were euthanized according to the experimental protocol or when humane endpoints were reached.

For CAR T cell detection by FACS in the bloodstream, 12.5 μl sample of peripheral blood was collected via tail vein bleeding and transferred to EDTA-coated tubes (MiniCollect K2E K2EDTA tubes, Greiner Bio-One) and treated for 5 min with ACK lysis buffer (Gibco). Cell suspensions were washed once with PBS and stained in 100 µl DPBS (Gibco) with 1 % FCS (Biowest). Antibodies were added using the antibody to cell ratio recommended by the supplier. Antibodies used are: CD45-BV510 (clone HI30, Biolegend), CD3-BV421 (clone UCHT1, Biolegend), CD4-PerCP/Cy5.5 (clone SK3, Biolegend), CD8α-APC/FIRE750 (clone SK1, Biolegend). After antibody incubation, cells were washed once with FACS buffer and propidium iodide (Invitrogen) was added. Flow cytometric analysis was performed on a BD FACS Symphony A3. Data was analyzed using FACS DIVA software (BD Biosciences) and FlowJo.

### Statistical analysis

Statistical analysis was caried out with GraphPad Prism v10.2.2. All *in vitro* data is presented as mean ± SD, unless otherwise noted in the figure legend. Statistical significance in RNA expression (Fig. 2A), CAR T cell - tumor organoid co-cultures (Fig. 3B, 3F, 4F-G, S3G) and LegendPlex experiments with multiple effector ratios (Fig. S3G) was determined by two-way analysis of variance (ANOVA) with Tukey’s multiple comparisons test, with a single pooled variance. Statistical significance in LegendPlex experiment with 1:10 effector ratios (Fig. 3D, 3G, S3H) was determined by two-way ANOVA with Šídák’s multiple comparisons test, with a single pooled variance. Significance was considered at P <0.05. P-values are indicated as *P < 0.05, **P < 0.01, ***P < 0.001, ****P < 0.0001. All *in vivo* data is presented as mean ± SEM, unless otherwise noted in the figure legend. Statistical significance in tumor volume graphs (Fig. 5B, S6A) was determined by two-way ANOVA with Turkey’s multiple comparison test at day 32 after CAR T cell injection. Statistical significance in Kaplan-Meier survival curves (Fig. 5C, S5B, S5E, S6B) is determined by log-rank (Mantel-Cox test). Statistical significance in CAR T cell detection by FACS (Fig. 5F, S6E) is determined by two-way ANOVA with Šídák’s multiple comparison test. Mice scarified due to reaching humane endpoints, retained the maximum value until the end of the experiments to avoid incorrect improvement of grouped tumor volumes due to dropouts. Significance was considered at P <0.05. P-values are indicated as *P < 0.05, **P < 0.01, ***P < 0.001, ****P < 0.0001.

## Data Availability

The data generated in this study are available within the article and its supplementary data files. Additionally, data can be accessed as follows:

*Cell surface proteomics:* Proteomics data including raw MaxQuant and data analysis files have been deposited to the ProteomeXchange Consortium via the PRIDE partner repository with the dataset identifier PXD046834 and data was used to generate Fig. 1C-D, Fig. S1A-D, and Suppl. Table 1.

*RNA-seq/snRNA-seq*: The processed bulk RNA sequencing data of ATRTs were obtained from the EGA database (EGAS00001006866: ATRT tissues n=7, ATRT organoids n=9) and for MRTs (EGAS00001003853: MRT tissues n=10, MRT organoids n=7, normal kidney organoids n=5) and used to for the analyses performed in Fig. 2A and Fig. 2C. The processed sequencing data of combined single-nuclei RNA and ATAC data (eMRT tissues n=7) were from the GEO database (accession code <u>GSE218385</u>) were used for the analyses performed in Fig. 2D and S2B).

*M&M dataset:* Gene expression data was obtained from the public Zenodo repository (https://doi.org/10.5281/zenodo.14167359) for the analyses performed in Fig. 4A-B, Fig. S4A-B.

## Acknowledgements

The work presented here is supported by KiKa core funding (to C.Y.J. and J.D.), Oncode InsÜtute, which is partly financed by the Dutch Cancer Society (to J.D), Oncode Accelerator, a Dutch NaÜonal Growth Fund project under grant number NGFOP2201 (to C.Y.J and J.D.), FoundaÜon Nikai 4 Life (to J.D), American AssociaÜon for Cancer Research (AACR) (to J.D), the St. Baldrick’s FoundaÜon (Pediatric Cancer Research Award to J.D.), SÜchÜng tegen Kanker (C/2024:2562 to B.V). J.L.B. received financial support from the Deutsche Forschungsgemeinschaft (#458375005), S.d.M. is funded by a personal FWO grant (12AP724N). We want to thank the Drost, Janda, Nierkens, Molenaar and Kemmeren groups, Prof. Dr. Max van Noesel, and the Pathology Department of the Princess Máxima Center for thoughtful discussion and helpful feedback, and the Princess Máxima Center Imaging and Flow Cytometry and Cell Sorting Core Facilities for their assistance with imaging and FACS experiments, and Stephanie Schubert for her valuable contribution in proof-reading the manuscript. In the final steps of the preparation of this manuscript the authors used Grammarly and ChatGPT to improve the language. After using these tools, the authors reviewed and edited the content as needed and take full responsibility for the content. We are profoundly grateful to the patients and parents who agreed to participate in our research.

## Author contributions

C.Y.J. and J.D. conceived the project, secured funding, and supervised the research. J.L.B. and J.K. performed cell surface biotinylation and sample preparation. D.J.K.C. and W.W. conducted mass spectrometry experiments, processed raw data, and performed proteomics analyses. J.L.B., J.K., K.N., and L.H. carried out validation experiments for target expression. V.C. and P.K. performed RNA expression analyses of targets across pediatric solid tumors, while I.P. analyzed target expression in single-nuclei RNA-seq and ATAC-seq datasets. N.v.H. and E.d.B. performed and interpreted IHC experiments. A.M.C, M.v.H., J.L.B. and S.N. generated CAR T cells and provided key guidance on CAR T cell *in vitro* and *in vivo* experiments. J.L.B., B.Z., and K.N. conducted *in vitro* co-culture assays and J.L.B, B.Z. and J.K. analyzed data. S.d.M., J.L.B., and B.V. performed and analyzed *in vivo* CAR T cell efficacy studies. C.Y.J., J.D., J.L.B., and J.K. wrote the manuscript, with input from all authors. All authors reviewed and approved the final manuscript.

