## Supplementary Figures 1-6 for "Cell surface proteomics of patient-derived malignant rhabdoid tumor organoids identifies ROBO1 as potential CAR T cell target for pediatric solid tumors"

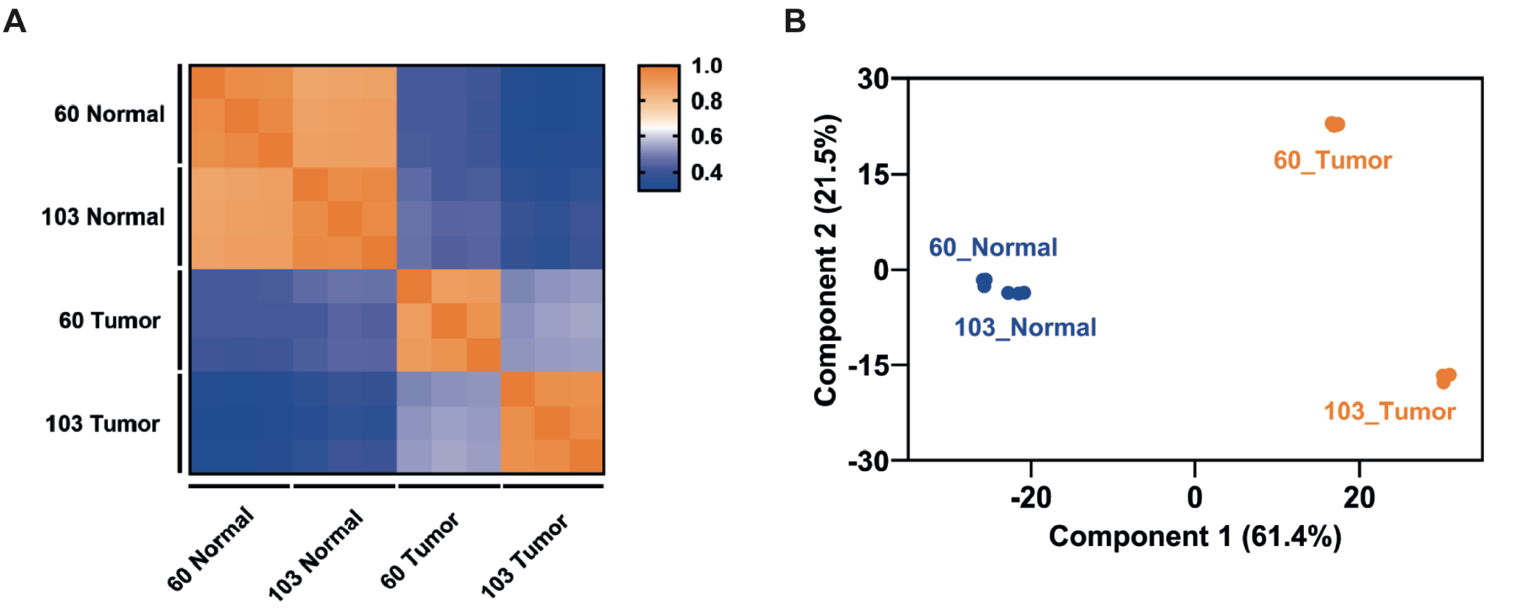


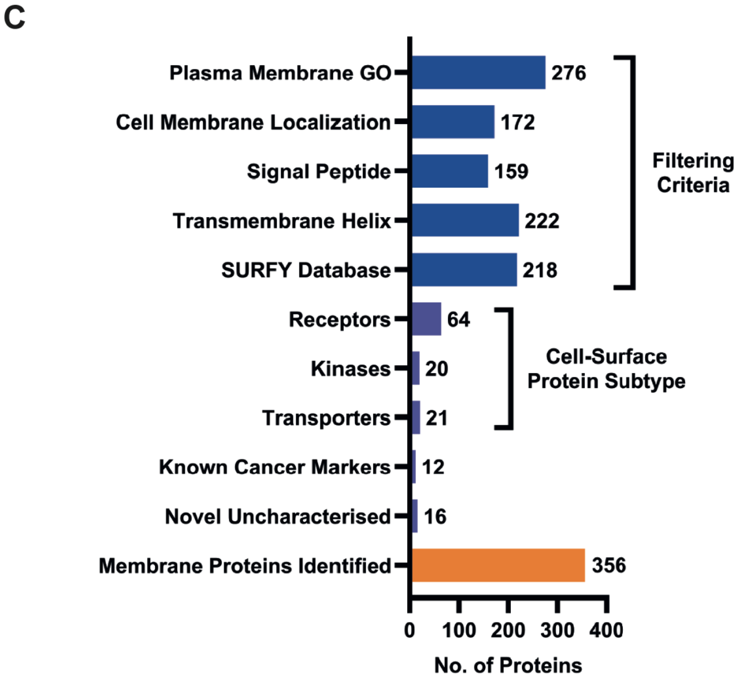


D

| Target | Fold Overexpression | |
|---|---|---|
| | 103T vs 103H | 60T vs 60H |
| ROBO1 | 118.6 | 2.7 |
| CSPG4 | 100.7 | 3.4 |
| MICB | 9.2 | 2.6 |
| TNFRSF10B | 5.8 | 11.1 |

**Supplementary Figure 1. Cell surface proteomics of patient-derived MRT organoids identifies potential targets for CAR T cell therapy.**
(A) Pearson correlation plot and (B) principal component analysis plot comparing the proteomes of two patient-matched normal kidney and eMRT organoids.
(C) Distribution of identified membrane proteins based on filtering criteria, cell surface protein subtype, known cancer-associated markers and novel uncharacterized proteins.
(D) Table showing fold overexpression of ROBO1, CSPG4, MICB and TNFRSF10B in 103T vs. 103H organoids and 60T vs. 60H organoids.

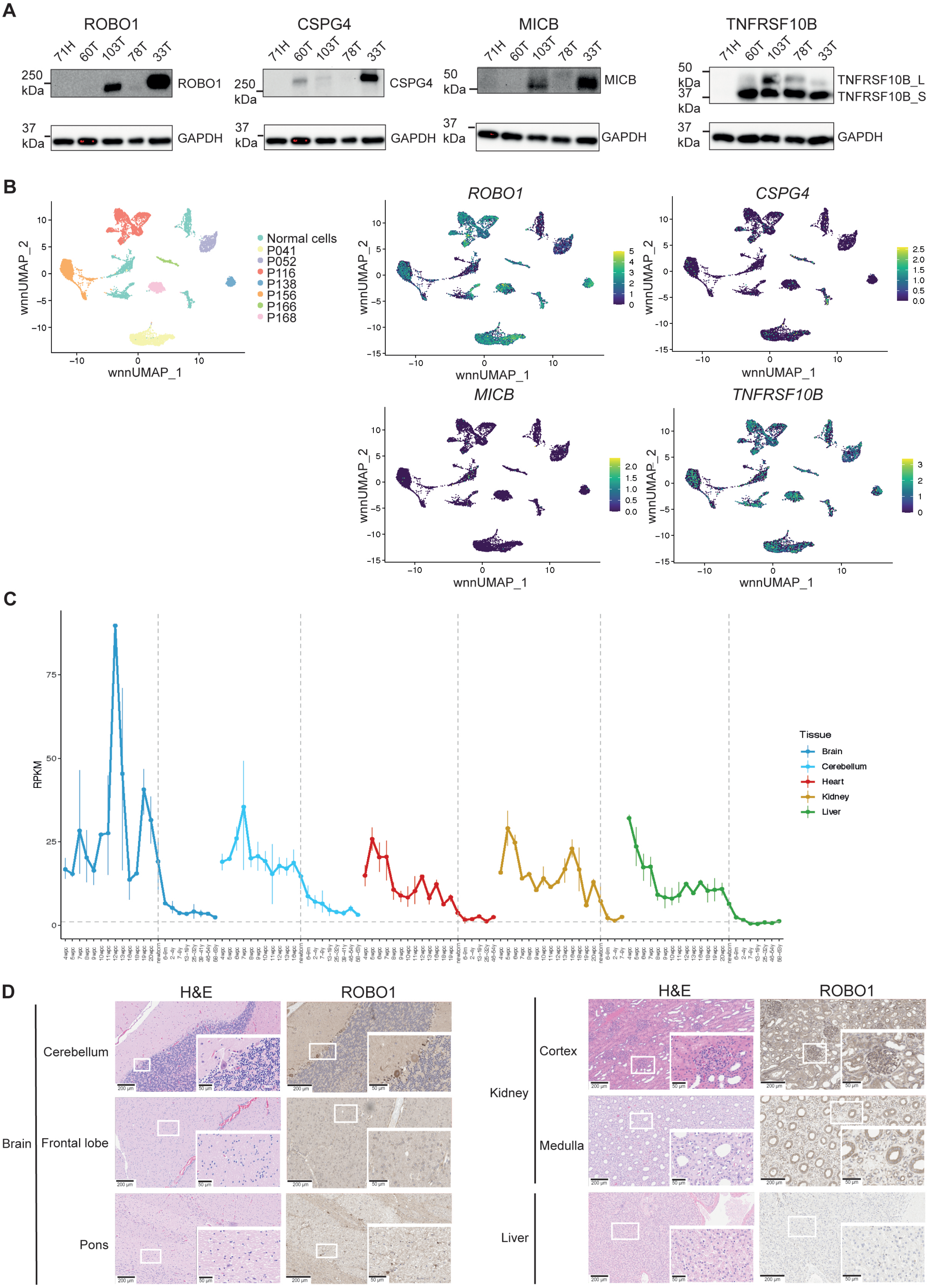

A
ROBO1
CSPG4
MICB
TNFRSF10B
71H 60T 103T 78T 33T
250 kDa
50 kDa
37 kDa
GAPDH
TNFRSF10B_L
TNFRSF10B_S
B
Normal cells
P041
P052
P116
P138
P156
P166
P168
wnnUMAP_1
wnnUMAP_2
ROBO1
CSPG4
MICB
TNFRSF10B
C
RPKM
Tissue
Brain
Cerebellum
Heart
Kidney
Liver
D
H&E
ROBO1
Brain
Cerebellum
Frontal lobe
Pons
Kidney
Cortex
Medulla
Liver

**Supplementary Figure 2. Gene and protein expression analysis validates ROBO1, CSPG4, MICB and TNFRSF10B upregulation in MRT organoids and tissues.**

(A) Western blot analysis of ROBO1, CSPG4, MICB, and TNFRSF10B in one normal kidney organoid model (H) and four eMRT organoids (T).

(B) UMAP visualization of main cell types in eMRT tissues, along with the expression patterns of *ROBO1*, *CSPG4*, *MICB* and *TNFRSF10B* genes.

(C) Temporal expression of *ROBO1* across normal tissues (brain, cerebellum, heart, kidney, and liver) during development. The dotted vertical line indicates the newborn stage.

(D) Representative IHC images of ROBO1 expression in normal brain (cerebellum, frontal lobe, and pons), kidney (cortex and medulla) and liver tissues.

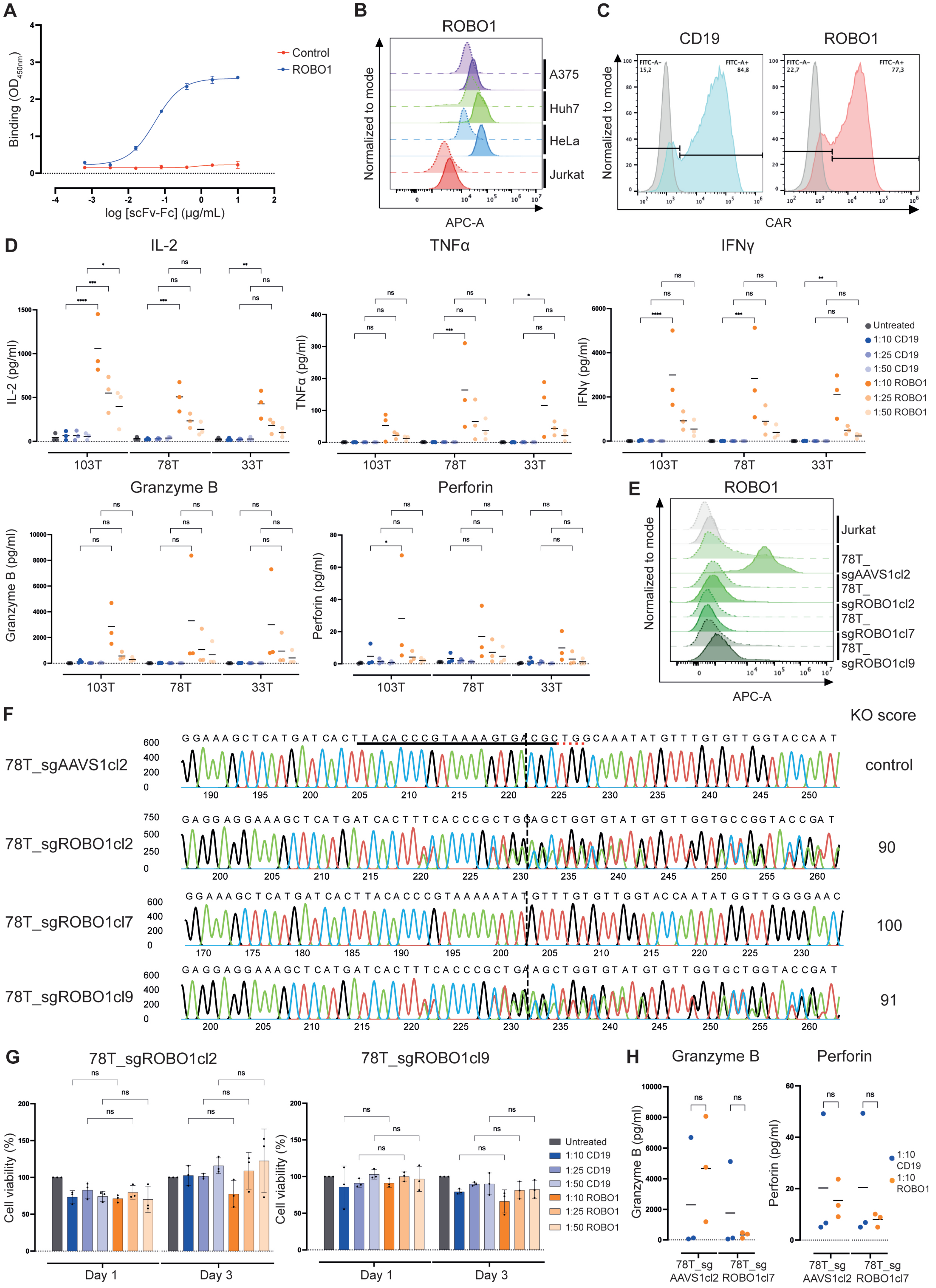

A
Binding (OD450nm)
log [scFv-Fc] (μg/mL)
Control
ROBO1
B
ROBO1
Normalized to mode
APC-A
A375
Huh7
HeLa
Jurkat
C
CD19
ROBO1
FITC-A-
15,2
FITC-A+
84,8
FITC-A-
22,7
FITC-A+
77,3
Normalized to mode
CAR
D
IL-2
TNFα
IFNγ
IL-2 (pg/ml)
TNFα (pg/ml)
IFNγ (pg/ml)
103T
78T
33T
Untreated
1:10 CD19
1:25 CD19
1:50 CD19
1:10 ROBO1
1:25 ROBO1
1:50 ROBO1
Granzyme B
Perforin
Granzyme B (pg/ml)
Perforin (pg/ml)
E
ROBO1
Normalized to mode
APC-A
Jurkat
78T_sgAAVS1cl2
78T_sgROBO1cl2
78T_sgROBO1cl7
78T_sgROBO1cl9
F
KO score
78T_sgAAVS1cl2
control
GGAAAGCTCATGATCACTTACACCCGTAAAAGTGACGCTGGCAAATATGTTTGTGTTGGTACCAAT
78T_sgROBO1cl2
90
GAGGAGGAAAGCTCATGATCACTTTCACCCGCTGGAGCTGGTGTATGTGTTGGTGCCGGTACCGAT
78T_sgROBO1cl7
100
GGAAAGCTCATGATCACTTACACCCGTAAAAATATGTTTGTGTTGGTACCAATATGGTTGGGGAAC
78T_sgROBO1cl9
91
GAGGAGGAAAGCTCATGATCACTTTCACCCGCTGAAGCTGGTGTATGTGTTGGTGCTGGTACCGAT
G
78T_sgROBO1cl2
78T_sgROBO1cl9
Cell viability (%)
Day 1
Day 3
Untreated
1:10 CD19
1:25 CD19
1:50 CD19
1:10 ROBO1
1:25 ROBO1
1:50 ROBO1
H
Granzyme B
Perforin
Granzyme B (pg/ml)
Perforin (pg/ml)
78T_sg AAVS1cl2
78T_sg ROBO1cl7
1:10 CD19
1:10 ROBO1

**Supplementary Figure 3. ROBO1-targeting CAR T cells elicit strong and specific anti-tumor activity in eMRT organoids.**

(A) ELISA and (B) flow cytometry confirming the binding of the recombinant anti-ROBO1 scFv-Fc to recombinant ROBO1 and cell surface-expressed ROBO1, respectively. Representative figures of three biological replicates. ELISA error bars represent SD of two technical replicates. Dashed lines in flow cytometry histograms indicate secondary antibody controls.

(C) Flow cytometry analysis of CAR expression on ROBO1 and CD19 CAR T cells.

(D) Cytokine production after 1-day co-culture of CD19 or ROBO1 CAR T cells with eMRT organoids at varying E:T ratios. Mean of three biological replicates, each containing the mean of two technical replicates.

(E) Flow cytometry histogram of ROBO1 expression on Jurkat cells, control (78T_sgAAVS1cl2) and three ROBO1 (78T_sgROBO1cl2, cl7 and cl9) knock-out eMRT organoid models. Solid lines: ROBO1 antibody staining; dashed lines: secondary controls. Representative data of three biological replicates.

(F) Sanger sequencing validation of ROBO1 knock-out (78T_sgROBO1cl2, cl7 and cl9) vs. AAVS1 control knock-out (78T_sgAAVS1cl2), highlighting sgRNA target and PAM sequence.

(G) Cell viability of ROBO1 knock-out eMRT organoids (78T_sgROBO1cl2 and cl9) after 1 or 3 days of co-culture with ROBO1 or CD19 CAR T cells. Results were normalized to untreated organoids. Mean ± SD of three biological replicates, each consisting of the mean of at least three technical replicates.

(H) Cytokine production after 1-day co-culture of CD19 or ROBO1 CAR T cells with control (78T_sgAAVS1cl2) or ROBO1 knock-out eMRT organoids (78T_sgROBO1cl7) at an E:T of 1:10. Mean of three biological replicates, each consisting of the mean of two technical replicates.

Statistical significance as in Fig. 2A.

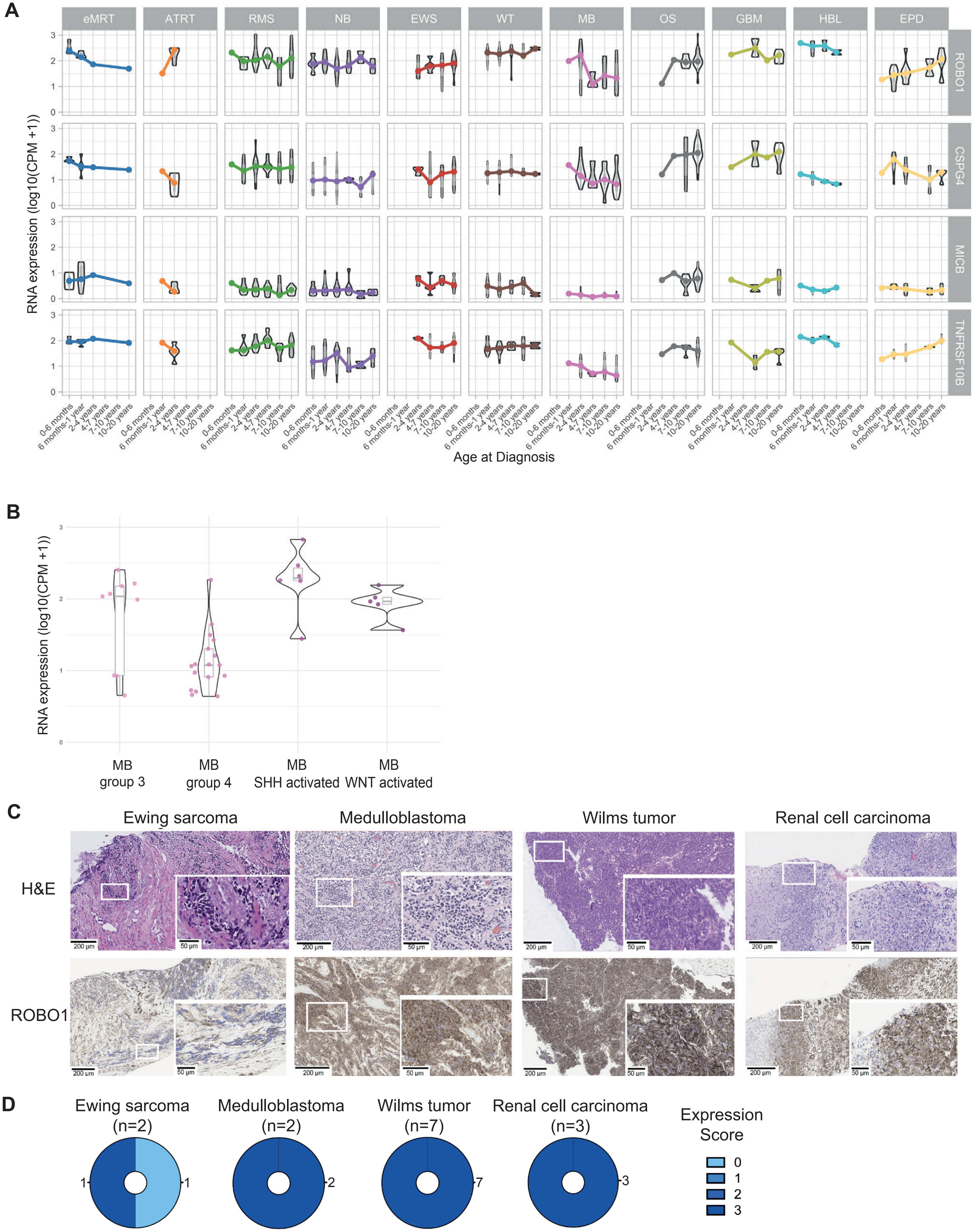
A
eMRT
ATRT
RMS
NB
EWS
WT
MB
OS
GBM
HBL
EPD
ROBO1
CSPG4
MICB
TNFRSF10B
RNA expression (log10(CPM +1))
0-6 months
6 months-1 year
2-4 years
4-7 years
7-10 years
10-20 years
Age at Diagnosis
B
RNA expression (log10(CPM +1))
MB
group 3
MB
group 4
MB
SHH activated
MB
WNT activated
C
Ewing sarcoma
Medulloblastoma
Wilms tumor
Renal cell carcinoma
H&E
ROBO1
200 µm
50 µm
D
Ewing sarcoma
(n=2)
Medulloblastoma
(n=2)
Wilms tumor
(n=7)
Renal cell carcinoma
(n=3)
Expression
Score
0
1
2
3

**Supplementary Figure 4. ROBO1 is a promising CAR T cell target for other pediatric solid tumors.**
(A) Violin plot showing RNA expression of *ROBO1, CSPG4, MICB, and TNFRSF10B* in pediatric solid tumors, stratified by age at diagnosis.
(B) Violin plot of *ROBO1* RNA expression in medulloblastoma, categorized by subtype.
(C) Representative IHC images of ROBO1 expression in Ewing sarcoma, medulloblastoma, Wilms tumor and renal cell carcinoma tissues, and expression scores (D).

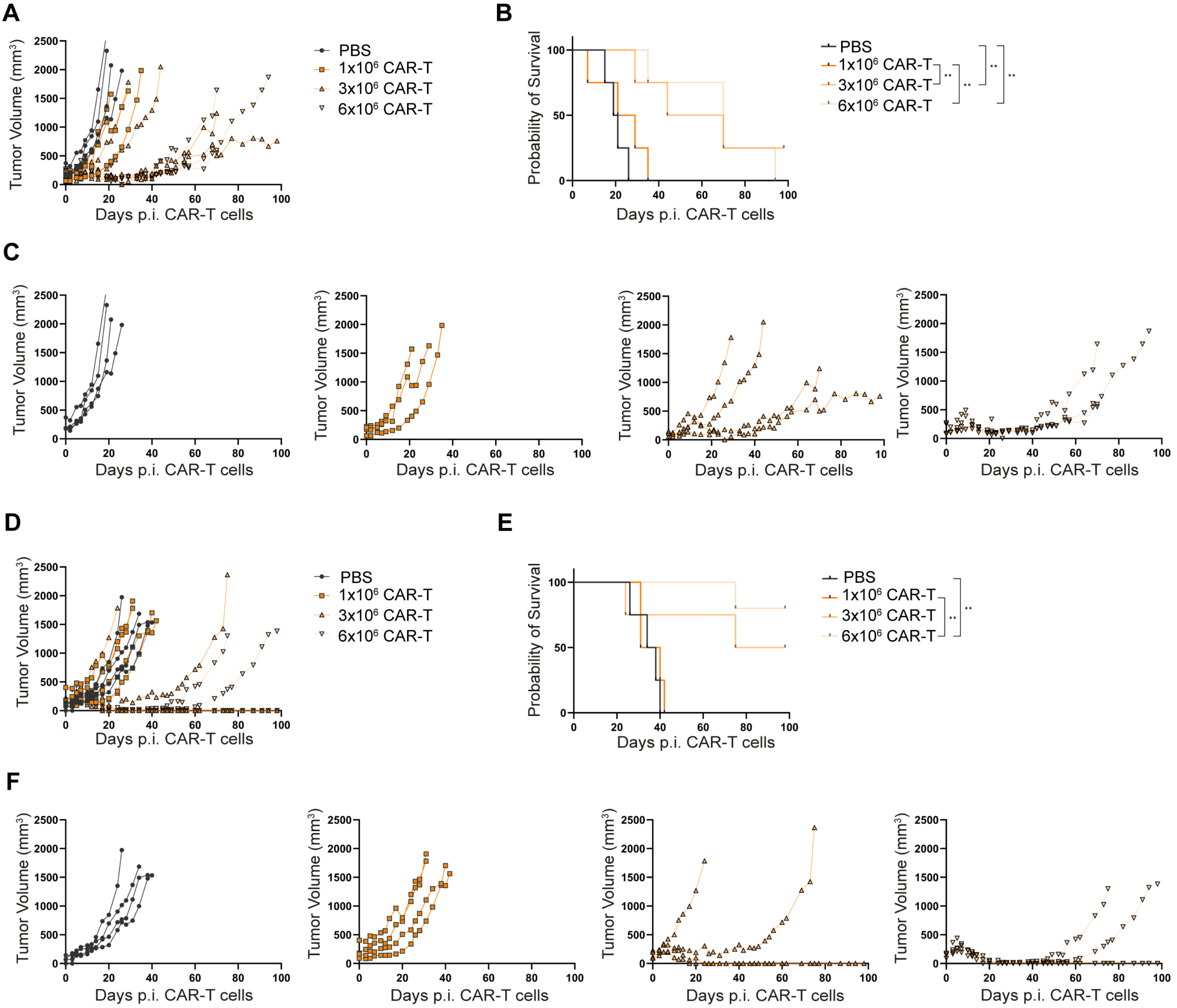
A
B
C
D
E
F
Tumor Volume (mm3)
Days p.i. CAR-T cells
Probability of Survival
PBS
1x10^6 CAR-T
3x10^6 CAR-T
6x10^6 CAR-T
**

**Supplementary Figure 5. ROBO1 CAR T cells exhibit dose-dependent functionality in eMRT PDX mouse models.**

(A) Tumor growth of 33T organoids following injections of PBS or increasing doses ROBO1 CAR T cells ($1x10^6$, $3x10^6$, and $6x10^6$), presented as the mean tumor volume ± SEM (n=4/group).

(B) Kaplan-Meier survival curves of mice in (A) treated with PBS or ROBO1 CAR T cells.

(C) Individual growth trajectories of tumors in mice in (A) treated with PBS or CD19 CAR T cells.

(D) Tumor growth of 103T organoids following injections of PBS or increasing doses ROBO1 CAR T cells ($1x10^6$, $3x10^6$, and $6x10^6$), presented as mean tumor volume ± SEM (n=4/group).

(E) Kaplan-Meier survival curves of mice in (D) treated with PBS or ROBO1 CAR T cells.

(F) Individual growth trajectories of tumors in mice in (D) treated with PBS or ROBO1 CAR T cells.

Statistical significance as in Fig. 2A.

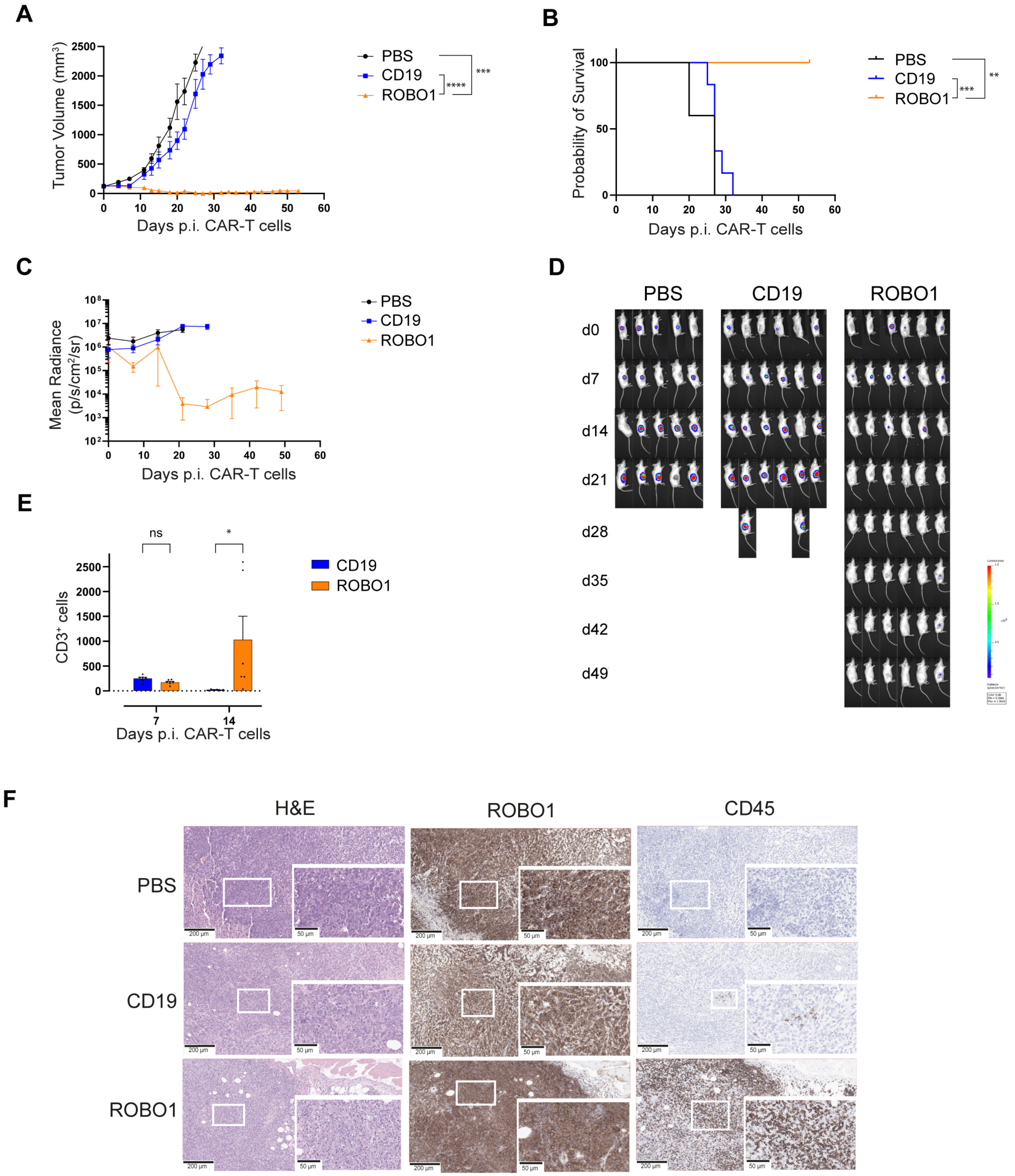

A
Tumor Volume (mm3)
Days p.i. CAR-T cells
PBS
CD19
ROBO1
***
****
B
Probability of Survival
Days p.i. CAR-T cells
PBS
CD19
ROBO1
**
***
C
Mean Radiance (p/s/cm2/sr)
Days p.i. CAR-T cells
PBS
CD19
ROBO1
D
PBS
CD19
ROBO1
d0
d7
d14
d21
d28
d35
d42
d49
E
ns
*
CD3+ cells
CD19
ROBO1
7
14
Days p.i. CAR-T cells
F
H&E
ROBO1
CD45
PBS
CD19
ROBO1
200 µm
50 µm

**Supplementary Figure 6. ROBO1-targeting CAR T cells induce potent and sustained cytotoxicity, eradicating eMRTs in vivo.**
(A) Growth of 33T-engrafted organoids in mice following CAR T cell injections, presented as mean tumor volume ± SEM (n=5 for PBS, n=6 for CD19 and ROBO1 CAR T cells).
(B) Kaplan-Meier survival curves of mice in (A).
(C) Longitudinal tumor burden from mice in (A) monitored by BLI.
(D) Representative BLI images of mice in (A).
(E) Flow cytometry analysis of peripheral blood from mice in (A), showing post-treatment immune cell counts.
(F) Representative IHC images tumor tissues from mice, assessing human ROBO1 and human CD45 expressing in treated mice.
Statistical significance as in Fig. 2A.
